# Lymphocyte alterations and elevated complement signalling are key features of refractory myasthenia gravis

**DOI:** 10.1101/2025.03.15.643341

**Authors:** Katherine C Dodd, Kirsten Baillie, James KL Holt, M Isabel Leite, Lijing Lin, Peter W West, James AL Miller, Jennifer Spillane, Stuart Viegas, Wioleta M Zelek, Jon Sussman, Madhvi Menon

## Abstract

**Background:** A significant proportion of patients with myasthenia gravis (MG) remain refractory to standard immunosuppressive therapy, and biomarkers to help guide treatment decisions are lacking.

**Methods:** We examined the circulating immune profile of patients with acetylcholine receptor antibody positive MG with differing treatment requirements and compared them to controls.

**Findings:** Refractory patients displayed the highest frequency of memory B cells, and increased production of interleukin (IL)-6 and tumour necrosis factor (TNF)-α upon toll-like receptor/ CD40 activation in vitro, mirrored by a dramatic loss of regulatory T cells (Tregs) and dendritic cells. Refractory MG was further characterised by elevated circulating complement proteins (C3, C5 and clusterin) and increased expression of complement receptors on lymphocytes. Following anti-CD20 therapy, residual plasmablasts persisted in circulation. Notably, a low baseline B cell frequency (<3%) was associated with poor clinical response to rituximab in refractory disease, although the sample size was limited.

**Conclusion:** Our findings define a distinct immune signature in refractory MG, identify potential biomarkers of treatment resistance, and highlight plasma cell depletion, IL-6 or complement inhibition, and Treg expansion as promising therapeutic avenues.

**Funding:** Funding was received from the NorthCare Charity, Myaware, Academy of Medical Sciences and the Neuromuscular Study Group.

**GRAPHICAL ABSTRACT:** 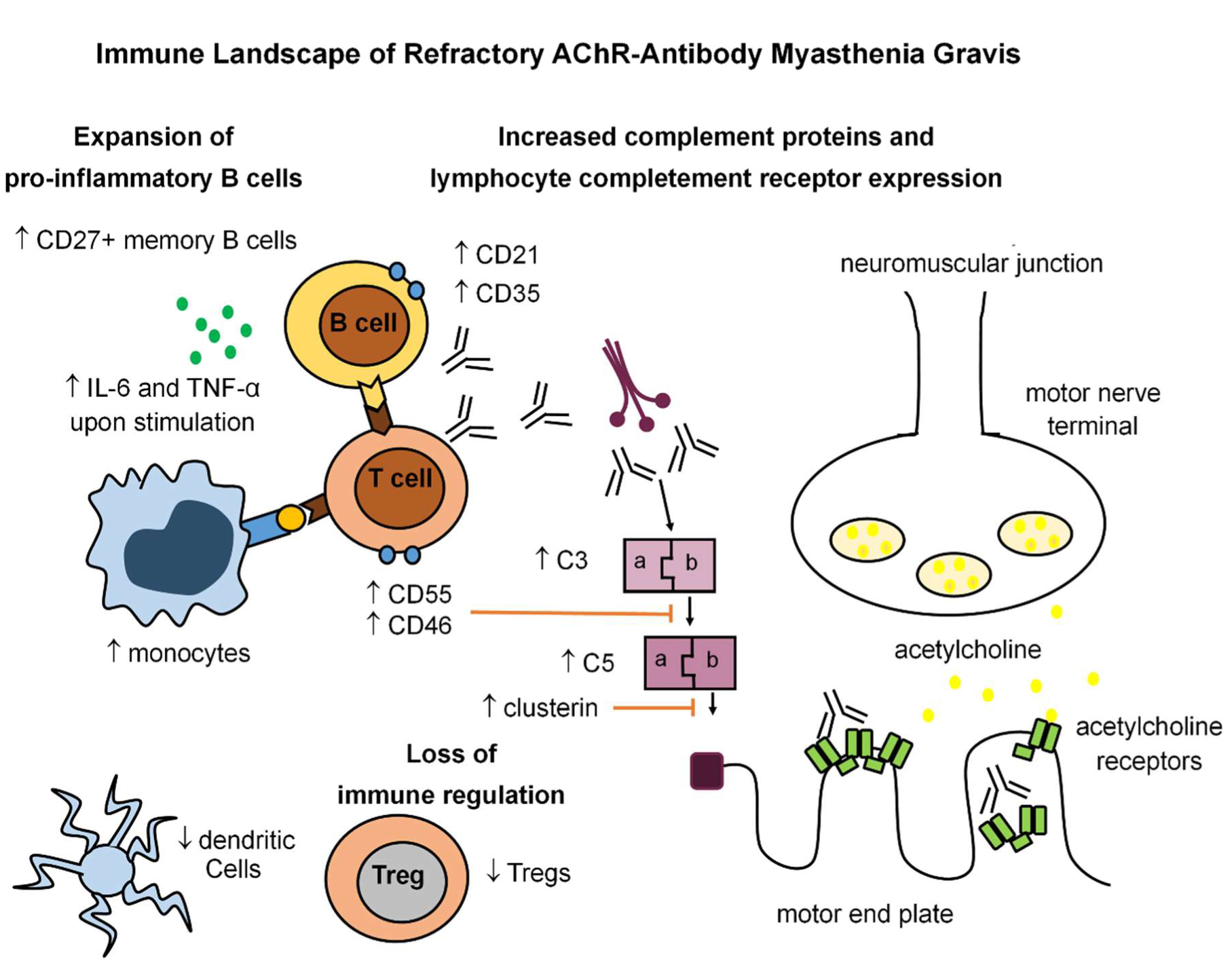

## INTRODUCTION

In Myasthenia Gravis (MG), autoantibodies target the neuromuscular junction, causing fatigable muscle weakness around the eyes (ocular) and often the bulbar, respiratory, and limb muscles (generalised). MG is a highly heterogenous disease of variable severity, with many refractory to standard immunosuppression, with resultant implications upon quality of life and large health-care resource utilization^1,2^.

Around 85% of cases are due to antibodies targeting the acetylcholine-receptor (AChR-ab)^3^. AChR-MG can be divided into three main sub-groups: (i) early-onset (EOMG), usually females in their 20-30s with thymic lymphoid follicular hyperplasia, (ii) late-onset (LOMG), usually males in their 60-70s, and (iii) thymoma-related. Generalised MG is initially treated with the acetylcholinesterase inhibitor pyridostigmine, followed by corticosteroids, and, if necessary, steroid-sparing immunosuppressants. B cell depleting agents have historically been reserved for refractory disease. Emerging therapies including complement, neonatal Fc receptor (FcRn) or IL-6 inhibitors show promise^4^, but patient selection is unclear.

There is a lack of reliable biomarkers to individualise treatment, and the pathophysiology of treatment resistance is poorly understood. AChR-ab titres do not correlate with disease severity, though the rate of change has been linked to outcome^5,6^. Clinical predictors of disease severity include thymoma, being overweight, and female gender^7,8^. MG is T cell-dependent^9^, with functional impairments in regulatory T cells (Tregs) previously reported^10,11^. Alterations in B cell populations have also been described, including an expansion of class-switched memory B cells, circulating plasmablasts and plasma cells^12–14^. Immunosuppression reduces naïve B cells and increases naïve T cells^12,15^, however, biomarkers indicative of an effective dose remain unknown. Previous immunophenotyping studies in MG have largely compared unstratified patient cohorts to healthy controls.

The innate immune system is increasingly recognised as contributing to autoimmune disease pathology. Myeloid cells, including dendritic cells (DC) and monocytes, are recognized to contribute to the initiation, perpetuation, and end-organ damage in MG^16–18^. Disruptions in DC and monocyte phenotypes have recently been implicated in MG pathophysiology^14,19,20^, however circulating myeloid cell frequencies remain poorly characterised.

AChR-abs activate the complement system, leading to membrane attack complex deposition at the neuromuscular junction^21^. Circulating complement findings in MG are inconsistent; some studies have reported lower circulating C3 and C4 levels^22,23^, whereas others have reported unchanged or elevated levels of C3, C3b and C5a^24,25^. Complement also modulates lymphocyte function, with C3 or C4 deficient experimental autoimmune MG (EAMG) mice exhibiting increased B cell apoptosis^26^. Complement receptor 2 (CD21) binds to C3d and promotes B cell survival, activation and antibody production^27,28^.It has been found to be upregulated on AChR-ab-producing B cells and correlated with AChR IgG titres^29^. In contrast, B cell complement receptor 1 (CD35), which binds to C3b/C4b and promotes regulatory B cell function, is downregulated on B cells in other autoimmune conditions^30,31^.

Complement regulators CD55 (decay-accelerating factor), CD59 (protectin), and CD46 (membrane cofactor protein) provide negative feedback to complement activation, and are expressed by T cells. EAMG mice lacking CD55 and/or CD59 are more susceptible to disease, and display a profoundly severe phenotype^32,33^. CD55, CD59 and CD46 also modulate T cell function^34,35^, however their expression on T cells in MG has not been previously characterised.

In the era of emerging high-cost targeted therapies for MG, identifying biomarkers that predict treatment resistance is of increasing clinical importance. Here, we undertook an observational cohort analysis to define immune cell profiles in refractory MG, with the aim of informing personalised therapeutic strategies and uncovering novel targets for intervention.

## RESULTS

### Clinical Cohorts

The study recruited 61 participants with AChR-MG and 21 controls, without autoimmune disease or cancer (solid organ or haematological). Participants with AChR-MG were stratified into four cohorts to distinguish between effects of disease severity and treatment: stable-non immunosuppressed (SNIS; stable on only low dose acetylcholinesterase inhibitors (≤120mg/day) for >2 years), stable immunosuppressed (SIS; stable for >2 years on azathioprine or mycophenolate mofetil with ≤5mg/day prednisolone), refractory (eligible for rituximab under NHS England criteria^36^; on corticosteroids and other immunosuppression with ongoing disease activity, or explosive bulbar onset), and treatment naïve (TN; diagnosed within the previous 6 months, no immunosuppression).

All participants with MG were AChR-ab positive at diagnosis; those with thymoma were excluded. The patient groups were compared to age- and gender-matched controls. Four participants were excluded (due to subsequent diagnosis of chronic myeloid leukaemia (n=1), thymoma (n=1), high CRP and suspicion of concurrent infection (n=1), or a grossly abnormal immune phenotype of uncertain cause (n=1)). The final cohort comprised 58 participants with AChR-MG (n=17 SNIS, n=23 SIS, n=10 refractory, n=8 TN) and 20 controls. Supplementary table 1 and Fig.1A show the demographics and clinical features of the included participants.

**Fig. 1.**
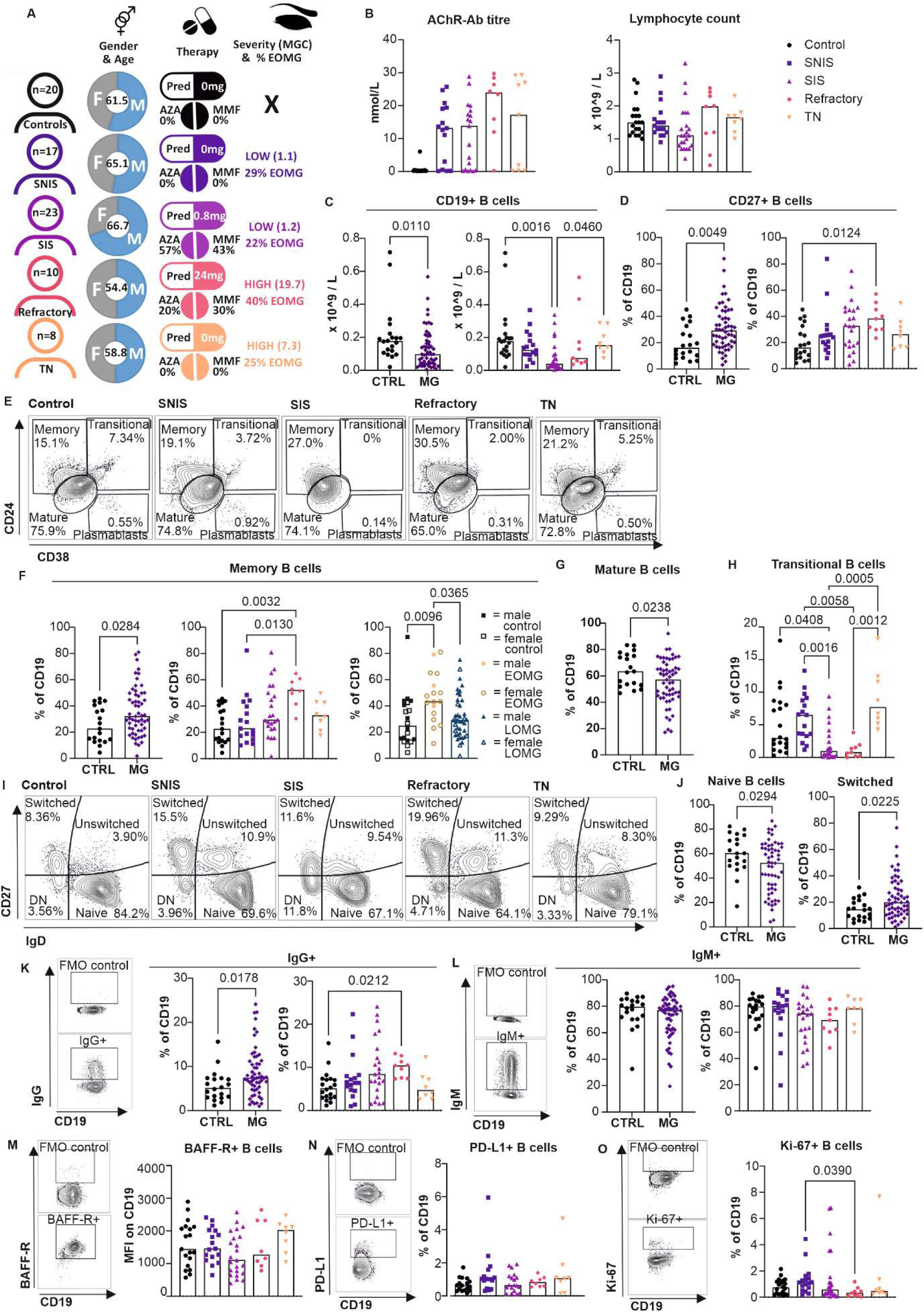
B cell alterations in patients with AChR-MG. **(A)** Infographic showing clinical characteristics, demographics and treatment of cohorts. **(B)** AChR titre in cohorts; control (n = 19), SNIS (n = 15), SIS (n = 19), refractory (n = 8), and TN (n = 9) and total lymphocyte count in cohorts; control (n = 20) SNIS (n = 17), SIS (n = 22), refractory (n = 8), and TN (n = 8). **(C)** CD19 count in analysis of PBMCs and **(D)** CD27^+^ frequency within CD19^+^ B cells, in control (n = 20) compared to MG (n = 57), and between subgroups; control (n = 20), SNIS (n = 17), SIS (n = 22), refractory (n = 9), and TN (n = 8). **(E)** Representative FACS plots for CD24 vs CD38, gated on CD19^+^ live lymphocytes, for control, SNIS, SIS, refractory and TN cohorts. **(F)** Memory B cell frequency within CD19^+^ B cells in control (n = 20) compared to MG (n = 57), between subgroups control (n = 20), SNIS (n = 17), SIS (n = 22), refractory (n = 9), and TN (n = 8), and between EOMG (n = 17) and LOMG (n = 40). **(G)** Mature B cell frequency within CD19^+^ B cells in control (n = 20) compared to MG (n = 57). **(H)** Transitional B cell frequency within CD19^+^ B cells in subgroups; control (n = 20), SNIS (n = 17), SIS (n = 22), refractory (n = 9), and TN (n = 8). **(I)** Representative FACS plots displaying CD27 and IgD expression on B cells, gated on CD19^+^ live lymphocytes, for control, SNIS, SIS, refractory and TN cohorts. **(J)** Switched and naïve (IgD^+^CD27^-^) B cell frequency within CD19^+^ B cells, in control (n = 20) compared to MG (n = 57). **(K)** IgG^+^ and **(L)** IgM^+^ B cell gating and frequency between control (n = 20) and MG (n = 57) and between subgroups; control (n = 20), SNIS (n = 17), SIS (n = 22), refractory (n = 9), and TN (n = 8). **(M)** BAFF-R gating strategy and mean fluorescent intensity on B cells, and **(N)** PD-L1^+^ and **(O)** Ki-67^+^ gating and B cell frequency in subgroups; control (n = 20), SNIS (n = 17), SIS (n = 22), refractory (n = 9), and TN (n = 8). Flow cytometry was performed on peripheral blood mononuclear cells (PBMCs). Graphs show individual participant data, with bar representing median values.

### Expansion of memory B cells in refractory MG

To determine whether B cell frequency better reflects effective immunosuppression than total lymphocyte count, peripheral blood mononuclear cells (PBMCs) were analysed by flow cytometry. AChR titre and total lymphocyte count did not differ between disease cohorts, and antibody titre did not correlate with disease severity (Fig.1B and S1A). We observed a significant reduction in CD19 count in MG compared to controls, primarily in the SIS cohort; but this did not distinguish between those on effective immunosuppression (SIS) and those refractory (Fig. 1C, gating strategy Fig. S1B). There was no difference in PBMC count between cohorts (Fig. S1C).

Further analysis revealed that CD27^+^ B cells were significantly expanded in MG, with the greatest increase in the refractory cohort when comparing disease subgroups (Fig. 1D and S1D-S1E). Next, we evaluated the expression of CD38 and CD24 on B cells and found that memory B cells were expanded in patients, again significantly in the refractory cohort (Fig. 1E-1F). When the patient cohort was grouped by age of disease onset, we observed an expansion of memory B cells and CD27^+^ B cells in those with EOMG compared to LOMG, independent of gender or age (Fig. 1F and S1E-S1F). Memory B cell frequency showed a weak positive correlation with disease severity (Fig. S1G).

Furthermore, we observed a reduction in mature B cell frequency in MG patients compared to controls (Fig. 1G), with no differences between MG subgroups (Fig. S1H). Both immunosuppressed groups (SIS and refractory) displayed a reduction in transitional B cell frequency (Fig. 1H), a known consequence of azathioprine and MMF^37^, with no overall difference between patients and controls (Fig. S1I). There were no differences observed in the frequency of circulating plasmablasts, when measured as CD38^++^CD24^-^, CD38^++^CD27^++^ or Blimp-1^+^ between cohorts (Fig. S1J-S1L).

We next characterised B cells into naïve, class-switched memory, unswitched memory, and double negative memory B cell subsets based on immunoglobulin D (IgD) and CD27 expression (Fig. 1I). MG patients showed an expansion of class-switched memory B cells accompanied by reduced naïve B cells compared to controls (Fig. 1J and S1M-S1N), with no significant differences in subset frequencies between EOMG and LOMG (Fig. S1N). IgG expression was increased in MG compared to controls (Fig. 1K), with no differences in IgM expression (Fig. 1L). Frequencies of unswitched and double negative memory B cells were comparable between patients and controls (Fig. S1O-S1P).

We examined the expression of BAFF-R and PD-L1 on B cells and did not observe any differences in relation to disease activity or treatment (Fig. 1M-1N and S1Q-S1R). Ki-67 expression (indicative of proliferation), was reduced in refractory MG compared to SNIS, but no significant differences were observed between other cohorts (Fig. 1O and S1R). This may be due to functional exhaustion of B cells in refractory disease or may reflect the more extensive immunosuppressive therapies used in refractory patients.

### B cells display a pro-inflammatory phenotype in refractory MG

Next, we stimulated PBMCs from MG patients and controls for 48 hours with CD40L, CpG-B (TLR9 agonist) or R848 (TLR7 agonist), and assessed cytokine production. Interestingly, MG patients produced significantly more pro-inflammatory cytokine IL-6 upon activation via TLR9, TLR7 or CD40 (compared to controls), and this was highest in those with refractory disease upon CD40 activation (Fig. 2A). TNF-α production was significantly increased in MG patients upon activation via TLR9 and TLR7, but not CD40 (Fig. 2B). When comparing disease subgroups, TNF-*α* production upon TLR9 activation was highest in the refractory cohort (Fig. 2B). IL-10 production was increased in MG patients only upon TLR9 activation, with no differences observed between patient subgroups (Fig. 2C). No differences were observed between those with EOMG and LOMG (Fig. 2A-2C).

**Fig. 2.**
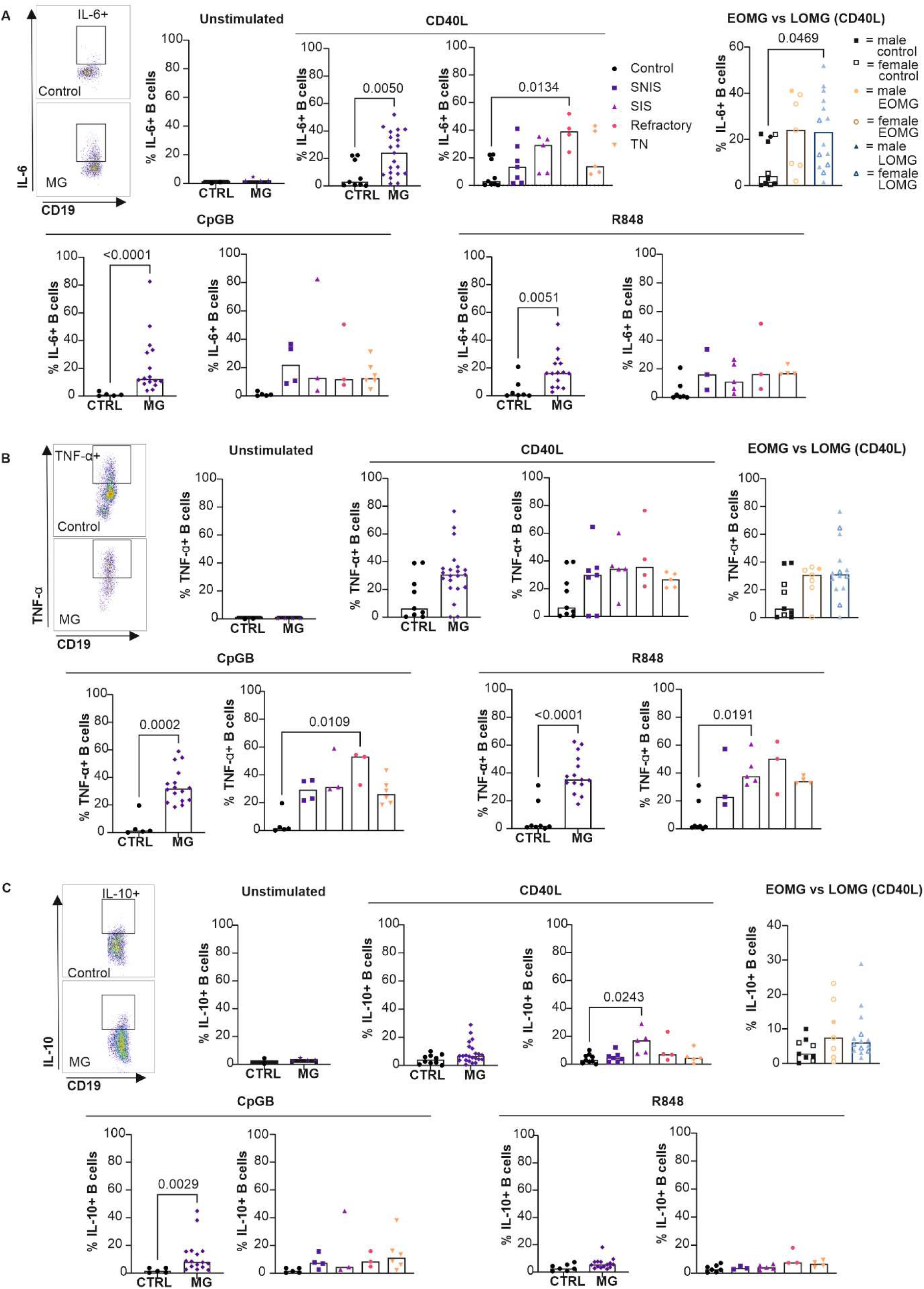
B cell cytokine production in patient with AChR-MG. Representative FACS plot demonstrating **(A)** IL-6^+^ ,**(B)** TNF-α^+^ and **(C)** IL-10^+^ B cells in control compared to MG and IL-6^+^ , TNF-α^+^ and IL-10^+^ B cell frequency when unstimulated and following stimulation with CD40L, CpG-B, and R848 in control (n = 5 CpG-B, n = 7 R848, n = 10 CD40L) compared to MG (n = 16 CpG-B, n = 15 R848, n = 23 CD40L), and when split into early-(n = 7) or late-onset MG (n = 15) Flow cytometry was performed on peripheral blood mononuclear cells (PBMCs). Graphs show individual participant data, with bar representing median values.

### Reduced frequencies of Tregs and dendritic cells in refractory MG

To determine if these B cell changes, or disease phenotype, may relate to changes in the T cell compartment, the T cell profile was analysed. CD3 count did not differ between groups (Fig. 3A), but CD4 frequency was higher in MG patients compared to controls, mainly in those on immunosuppression (Fig. 3B and S2A-S2B). CD8^+^ T cell frequencies did not significantly differ (Fig. S3C), but CD4^+^CD8^+^ double positive T cells were lower in MG, particularly in the SIS cohort (Fig. S3D).

**Fig 3.**
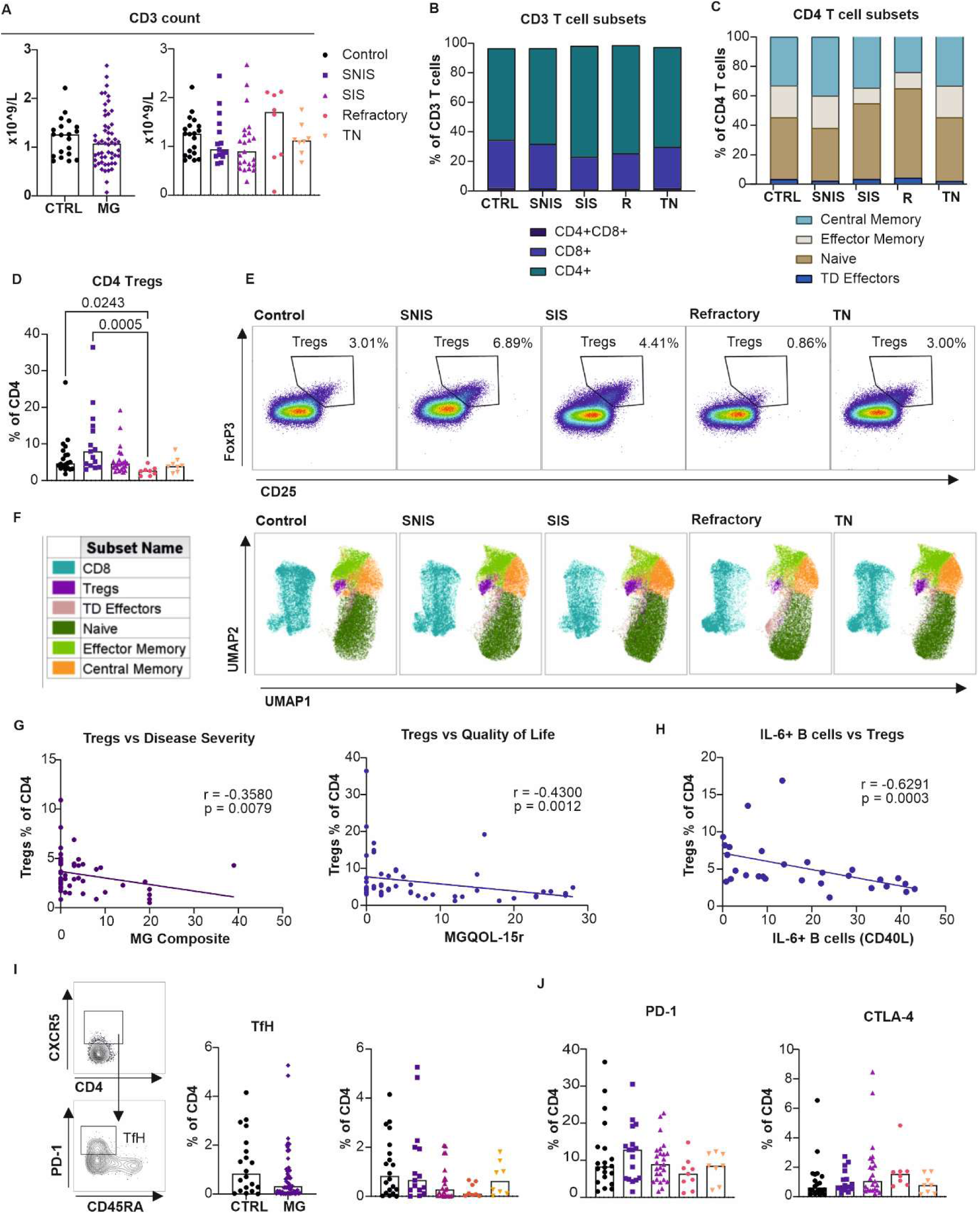
T cell alterations in patients with AChR-MG. **(A)** CD3 count, gated on live lymphocytes in PBMCs, in control (n = 20) compared to MG (n = 54), and between subgroups; control (n = 20), SNIS (n = 15), SIS (n = 23), refractory (n = 8), and TN (n = 8). **(B)** Proportion of CD4^+^, CD8^+^ and CD4^+^CD8^+^ cells, within CD3^+^ live lymphocytes, and **(C)** naïve, central and effector memory, and TD effector T cell frequency, gated on CD4^+^ T cells, in control (n = 20), SNIS (n = 15), SIS (n = 23), refractory (n = 8), and TN (n = 8). **(D)** FoxP3^+^CD25^+^ Treg frequency, gated on both CD4^+^ and CD8^+^ T cells, in control (n = 20), SNIS (n = 15), SIS (n = 23), refractory (n = 8), and TN (n = 8). **(E)** Representative FACS plots for Tregs in control, SNIS, SIS, refractory and TN cohorts. **(F)** UMAP plots demonstrating main T cell subsets in representative selection of 5,000 CD3^+^ live lymphocytes of control (n = 8), SNIS (n = 8), SIS (n = 8), refractory (n = 8) and TN (n = 8). **(G)** Simple linear regression for CD4 Treg frequency against MG Compositive and MG QoL-15r score (n = 54) **(H)** Simple linear regression for CD4 Treg frequency against IL-6^+^ B cells when stimulated with CD40L (n = 29). **(I)** FACS plot demonstrating gating of CXCR5^+^CD45RA^-^PD-1^+^ Tfh cells, gated on CD4^+^ T cells, and graphs showing CXCR5^+^CD45RA^-^PD-1^+^ Tfh cell frequency count in control (n = 20) compared to MG (n = 54), and between subgroups; SNIS (n = 15), SIS (n = 23), refractory (n = 8), and TN (n = 8). **(J)** PD-1+ and CTLA-4^+^ CD4 T cell frequencies in control (n = 20), SNIS (n = 15), SIS (n = 23), refractory (n = 8), and TN (n = 8). Flow cytometry was performed on peripheral blood mononuclear cells (PBMCs). Graphs show individual participant data, with bar representing median values.

Analysis of CD4^+^ T cell subsets revealed an expansion of naïve T cells in refractory MG (Fig. 3C and S2E-S2F). Effector memory T cells were reduced in both immunosuppressed groups (SIS and refractory), whereas central memory and TD effector cell frequencies did not significantly differ between subgroups (Fig. 3C and S2E-S2F). Notably, CD4^+^CD25^+^FoxP3^+^ Tregs were significantly reduced in the refractory cohort compared to controls and SNIS patients (Fig. 3D-3F). Treg frequencies negatively correlated with disease severity and disease-related quality of life scores (Fig. 3G), suggesting a potential role in ameliorating disease. No significant difference in CD8 Tregs was seen (Fig. S3G). As Tregs are known to suppress pro-inflammatory B cells, we next examined whether the loss of Tregs was associated with the expansion of effector B cells. Indeed, we observed a significant negative correlation between IL-6-producing B cells and Treg frequencies (Fig. 3H). No differences in Tregs were observed between EOMG and LOMG (Fig. S2H). In contrast to previous reports of expanded circulating Tfh cell frequencies in MG^38,39^, we observed a trend towards reduction in MG patients, particularly in the refractory cohort (Fig. 3I and S2I).

Further, we examined the expression of immune checkpoint molecules, PD-1 and cytotoxic T lymphocyte antigen 4 (CTLA-4), given that blockade of these pathways can induce MG-like syndromes and that genetic studies have linked CTLA-4 variants to MG^40,41–43^. PD-1 and CTLA-4 expression by CD4^+^ and CD8^+^ T cells, as well as Tregs did not significantly differ between MG cohorts and controls (Fig. 3J and S2J-S2K).

### Myeloid compartment changes seen in refractory-MG

Given the interplay between the adaptive and innate immune systems, we next investigated whether alterations in myeloid cells might contribute to reduced Treg frequencies in refractory MG (gating strategy in Fig. S3A). Although there were no differences in DC frequencies between all MG patients and controls, there was a dramatic reduction in the refractory MG cohort when comparing disease subgroups (Fig. 4A). Both pDC and mDC frequencies were decreased in the refractory cohort (Fig. 4B and S3B), however, the pDC:mDC ratios were similar between patient groups (Fig. S3C). Treg frequency does not correlate with DC frequency, but shows a weak correlation with pDC frequency (Fig. 4C). Of note, pDCs showed greater correlation with disease severity and quality of life scores than total DC frequency (Fig. 4C). Monocytes were expanded across MG cohorts (apart from SNIS) and skewed towards the classical phenotype (Fig. 4D-4E and S3D). NK cell frequencies and subsets did not show any significant differences between MG patients and control, or between patient cohorts (Fig. 4F-4G and S3E-S3F). The SIS group exhibited reduced frequencies of CD86^+^ monocytes and DCs, whereas no differences were observed in CD86 expression on NK cells and CD80 expression across all myeloid subsets (Fig. S3G-S3I).

**Fig. 4.**
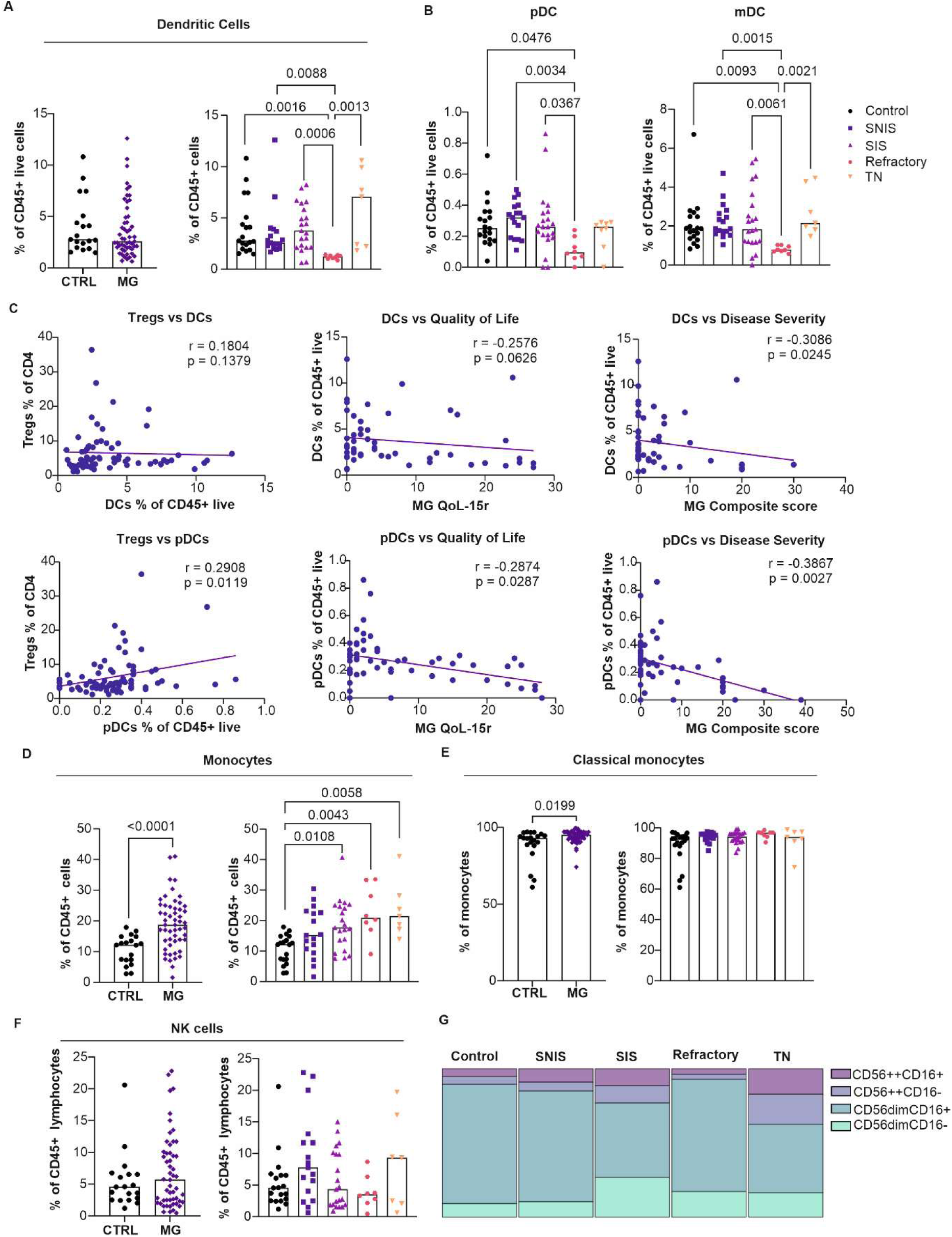
Myeloid cell alterations in patients with AChR-MG. **(A)** Frequencies of dendritic cells (CD3^-^CD19^-^CD66b^-^CD14^-^HLA-DR^+^) in control (n = 18) compared to MG (n = 48) and in subgroups; SNIS (n = 17), SIS (n = 19), refractory (n = 6), TN (n = 6). **(B)** pDC and mDC frequency in subgroups; control (n = 18), SNIS (n = 17), SIS (n = 19), refractory (n = 6), TN (n = 6). **(C)** Simple linear regression for DCs and pDCs against Tregs frequency (n = 74), quality of life (n = 53), and disease severity (n = 53). **(D)** Frequencies of monocytes (CD3^-^CD19^-^CD66b^-^CD14^+^HLA-DR^+^CD64^+^) in control (n = 18) compared to MG (n = 48) and in subgroups; SNIS (n = 17), SIS (n = 19), refractory (n = 6), TN (n = 6). **(E)** Frequencies of classical monocytes in control (n = 18), compared to MG (n = 48), and in subgroups; SNIS (n = 17), SIS (n = 19), refractory (n = 6), and TN (n = 6). **(F)** Frequencies of NK cells in control (n = 18) compared to MG (n = 48) and in subgroups; SNIS (n = 17), SIS (n = 19), refractory (n = 6), TN (n = 6). **(G)** NK cell subtypes in subgroups; control (n = 18), SNIS (n = 17), SIS (n = 19), refractory (n = 6), TN (n = 6). Flow cytometry was performed on peripheral blood mononuclear cells (PBMCs). Graphs show individual participant data, with bar representing median values.

### Complement receptor expression on lymphocytes and circulating complement proteins are elevated in refractory MG

Given that AChR-abs activate the complement system^44^, we examined circulating complement proteins (proteins TCC, Ba, C1q, C4, C9, CR1, FH, FHR4, FI, iC3b, properdin, and FHR125) in MG and its association with disease and treatment. Although there were no differences observed when comparing all MG patients to controls (Fig. S4A), or between EOMG and LOMG (Fig. S4B), the circulating levels of C3, C5 and clusterin were significantly elevated in refractory disease when comparing patient subgroups (Fig. 5A-5B and S4C). Further, C3, C5 and clusterin levels positively correlated with disease severity and quality of life scores (Fig. 5B, S4D).

**Fig 5.**
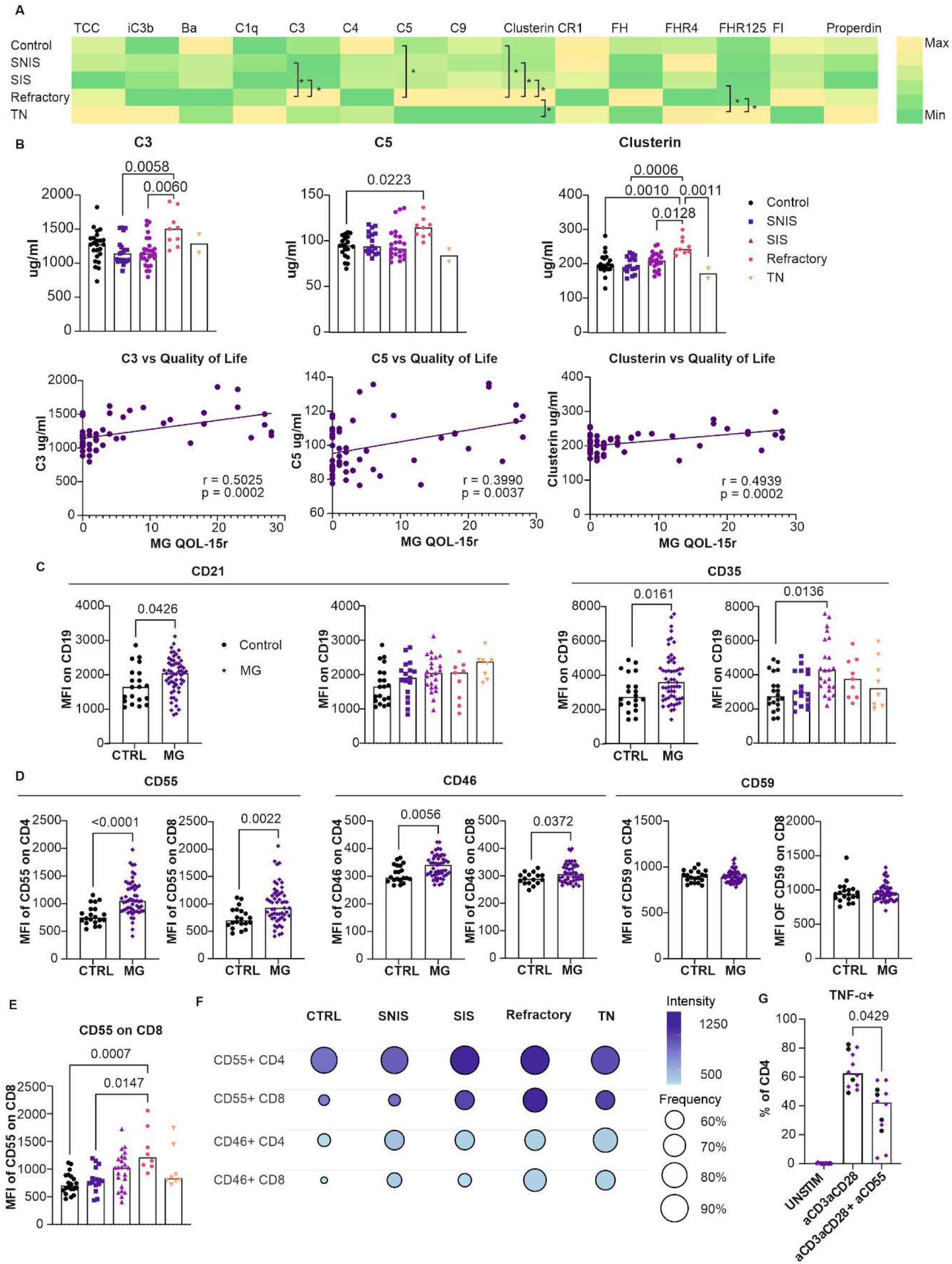
Elevated circulating complement proteins and expression of complement receptors on lymphocytes in refractory MG. Complement proteins were measured by ELISA and their receptor expression on PBMCs analysed by flow cytometry. **(A)** Heatmap demonstrating relative concentration of each complement protein analysed and **(B)** frequencies of C3, C5, and clusterin in subgroups; control (n = 20), SNIS (n = 18), SIS (n = 23), Refractory (n = 9), and TN (n = 2), and correlation between C3, C5, clusterin and quality of life (MG Qol-15r score) (n = 52). **(C)** Expression of CD21 and CD35 on B cells in control (n = 20) compared to MG (n = 56), and between subgroups; control (n = 20), SNIS (n = 17), SIS (n = 22), refractory (n = 9), and TN (n = 8). **(D)** Graphs showing expression of CD55, CD46 and CD59 on CD4 and CD8 T cells in control (n = 20), compared to MG (n = 54). **(E)** CD55 expression on CD8 T cells in subgroups; control (n = 20), SNIS (n = 15), SIS (n = 23), refractory (n = 8), and TN (n = 8). **(F)** Bubble plot demonstrating frequency and intensity of expression of CD46 and CD55 on CD4 and CD8 T cells in subgroups control (n = 20), SNIS (n = 15), SIS (n = 23), refractory (n = 8), and TN (n = 8). **(G)** TNF-α+ CD4 T cells in controls (n = 4) and MG (n = 8) when unstimulated, stimulated with aCD3aCD28 and when stimulated with anti-CD55. Graphs show individual participant data, with bar representing median values.

Complement is known to cause disruption at the neuromuscular junction^26^, however, its interaction with lymphocytes remains unclear. To explore this, we examined CD21 and CD35 expression on B cells, and discovered increased mean fluorescence intensity of both receptors in MG patients compared to controls (Fig. 5C). However, frequencies of CD35-expressing B cells, but not CD21-expressing B cells, were reduced in MG patients compared to controls, with no differences between patient subgroups (Fig. S4E).

We next examined the expression of complement receptors CD55, CD46, and CD59 on T cells (Fig. S4F-S4G). Intensity of expression of CD55 and CD46, but not CD59, was significantly higher on CD4 and CD8 T cells (Fig. 5D). When examined by treatment subgroups, CD55 expression was highest in the refractory cohort on CD8 T cells, and in both refractory and SIS cohorts on CD4 T cells (Fig. 5E-5F, and S4I), however, there was no significant difference in intensity of CD46 expression between subgroups (Fig S4H). Frequencies of CD55^+^ CD4/CD8 T cells and CD46^+^ CD4 T cells were significantly expanded in MG, with CD55^+^ and CD46^+^ CD8 T cell frequencies highest in the refractory cohort (Fig. 5F and S4J), indicating ongoing complement-mediated pathology in refractory disease. No differences were observed in CD59-expressing T cells.

Further, we stimulated lymphocytes with R848 and, anti-CD3 and anti-CD28, whilst inhibiting the complement receptors CD21, CD35 and CD55 to see if this impacted cytokine production in a subset of samples. There was significantly reduced production of TNF-α from CD4 T cells upon stimulation in the context of CD55 inhibition (Fig. 5G); no other significant differences were seen in the production of IL-6, IL-10, IL-17, TNF- α or IFN-γ (Fig. S5A-S5B).

### Unsupervised Cluster Analysis Reveals Defining Features of Refractory Disease

To determine whether a distinct immune signature characterizes refractory MG, we applied principal component analysis (PCA) followed by unsupervised K-means clustering to our flow cytometry dataset. This approach identified six discrete immune clusters (Fig. 6A). Notably, Cluster 1 was enriched for refractory MG patients and was distinguished by a set of 16 immune markers (Fig. 6B). Specifically, this refractory patient-enriched cluster exhibited increased frequencies of CD27⁺ B cells, memory and unswitched B cells, naïve T cells, and CTLA-4⁺ T cells, along with elevated CD55 expression on T cells. Cluster 1 also showed reduced frequencies of Tregs, central and effector memory T cells, CD80⁺ DCs, and both mature and Ki-67⁺ B cells. The patient-level heatmap illustrates that these immune alterations are consistently observed across most individuals in Cluster 1 (Fig. 6C), whereas the remaining clusters display more heterogeneous immune profiles. These findings support the existence of a distinct immune phenotype associated with refractory MG.

**Figure 6.**
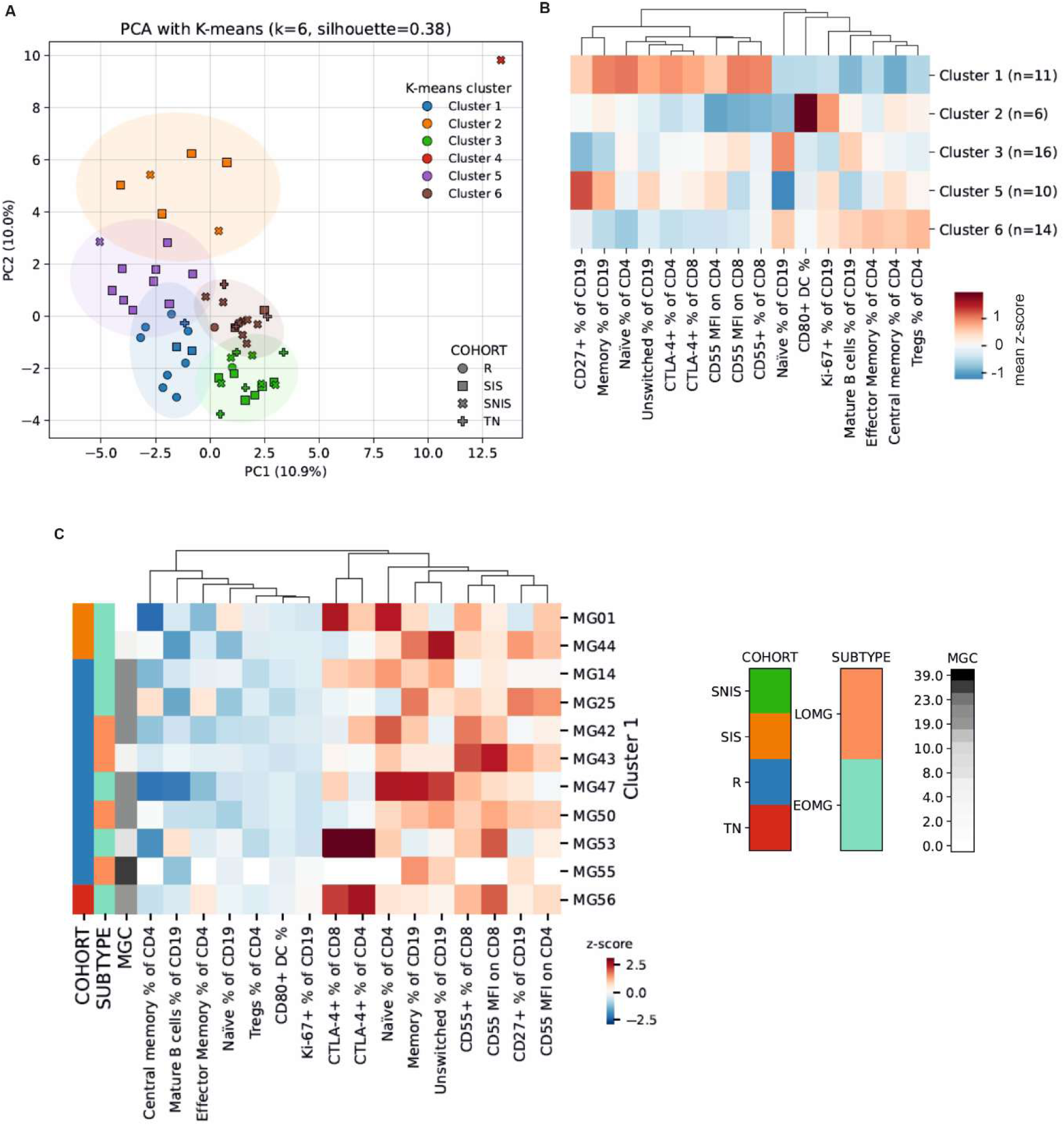
Distinct immune profile associated with refractory MG. (**A**) Two-dimensional principal component analysis of z-scored flow markers. K-means on [PC1, PC2] identified sic clusters. Points are colours by cluster and shaped by cohort; ellipses show 95% confidence contours in PC space. PC1 and PC2 explain 10.9% and 10.0% of variance, respectively. **(B)** Heatmap of mean z-scores for the top 16 varying markers across the six K-means clusters. Colours indicate mean z-scores (red = higher than the study-wide mean; blue = lower), centred at 0. **(C)** Heatmap showing cluster 1 per-sample pattern across the top discriminating markers.

### Low baseline B cell frequency is associated with poor response to rituximab

Eight of the 10 refractory participants were followed up at 1-, 7- and 13-months following B cell depletion with two infusions of rituximab (or biosimilar) 1 gram intravenously, two weeks apart. Some of these participants were given additional doses of rituximab at the clinician’s discretion; the timing of these doses, and changes in participant disease severity, quality of life, and prednisolone dose can be seen in Fig. 7A. No changes in AChR-ab titres were observed (Fig. S6A). All participants showed successful B cell depletion, with some participants displaying partial repopulation by month 13 (Fig. 7B-7C). Notably, those who did not reduce their severity score by 50% or more (defined as non-responders; R2, R4 and R7) had baseline B cell frequencies under 3%, whereas responders had baseline frequencies above 3%. Non-responders also had longer disease duration (median of 15 years), compared to responders (1 year).

**Fig. 7.**
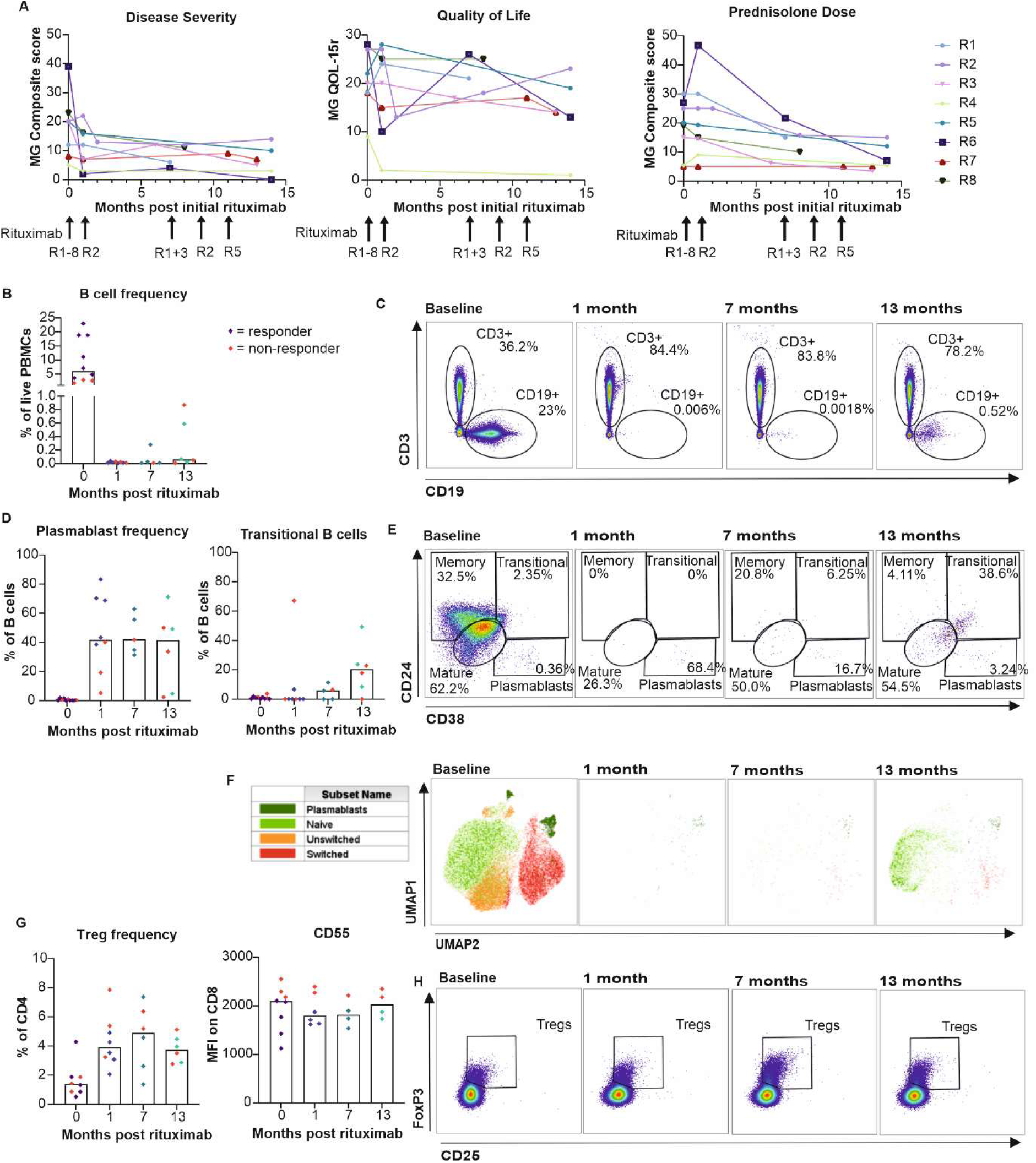
Immune alterations following B cell depletion therapy for refractory MG. **(A)** Disease severity, quality of life and prednisolone dose changes following rituximab therapy. Timing of additional doses indicated by arrows. Flow cytometry was performed on peripheral blood mononuclear cells (PBMCs) at baseline (n = 10), 1 month (n = 8), 7 months (n = 5), and 13 months (n = 6) post rituximab in the refractory group. **(B)** Changes in B cell frequency. **(C)** Representative FACS plots showing changes in B and T cell frequency. **(D)** Plasmablast and transitional B cell frequency. **(E)** Representative FACS plots showing changes in B cell subsets. **(F)** UMAP plots displaying main B cell populations in all CD19^+^ live lymphocytes (total=195,944 cells). **(G)** Changes in Treg frequency and CD55 expression on CD8 T cells. **(H)** Representative FACS plots showing changes in Treg frequencies, gated on CD4^+^ T cells. All graphs show individual participant data, with bar representing median values.

Of the remaining B lineage cells in circulation during depletion, a high proportion of these were found to be plasmablasts, alongside increased Blimp-1 expression (Fig. 7D-7E and S6A).

In agreement with previous findings^45^, the repopulating B cells (in months 7-13) were found to be transitional (CD24^++^CD38^++^) and mature (CD24^+^CD38^+^) B cells (Fig. 7D-7F). Residual B cells following depletion expressed less BAFF-R, CD21 and CD35, but not CD27 (Fig. S6A-S6B).

Treg frequencies increased following B cell depletion in both responders and non-responders (Fig. 7G-7H). There were no significant changes in CD4 or CD8 T cell frequencies, T cell phenotype, or expression of PD-1 or CD55 (Fig. S6C-S6E). It is noteworthy to mention that the rituximab clinical non-responders had the highest expression of CD55 on CD8 T cells at all time-points (before and after rituximab treatment), suggesting that there was ongoing complement-mediated pathology (Fig. 7G). Levels of C3, C5 and clusterin were unaltered in response to rituximab (Fig. S6F-S6G). In the myeloid compartment, there was no change to DC or monocyte frequencies, but a trend to increasing NK cell frequencies (Fig. S6H).

## DISCUSSION

In MG, a proportion of patients remain refractory to therapy, and biomarkers to predict disease course and thus to guide treatment decisions are lacking. This study is the broadest immunophenotyping study in AChR-MG to date, exploring circulating immune profiles in those with differing treatment requirements against age- and gender-matched controls. The pre-specified cohorts elucidated differences between those not requiring immunosuppression to obtain disease remission (SNIS), those responding to standard immunosuppression (SIS) and those who have not responded (refractory). We have identified a number of immune alterations that characterise refractory disease.

Circulating B cell frequencies are known to reduce with immunosuppression, or longstanding autoimmune disease^12^, a finding we replicate here. We also replicate others findings of class-switched memory B cells expansion in AChR-MG^14,46^, but show that this is most marked in refractory disease. We also identified that the expansion of memory B cells is in EOMG rather than LOMG, which may translate to differences in response to B cell-targeted therapies.

Surprisingly, we observed no differences in circulating plasmablast frequencies. This may be because AChR-abs are thought to be mainly produced by long-lived plasma cells, which reside in the bone marrow or lymphoid tissue rather than in circulation^44,47^. Two other studies have reported elevated plasmablast/plasma cell frequency in AChR-MG^12,13^, though these studies used different markers than this study to define these cell types and one did not include those on immunosuppression.

Although the pathogenic mechanism of MG is primarily effects of AChR-ab at the neuromuscular junction, we found that there is a broad circulating pro-inflammatory phenotype, with B cells primed to produce pro-inflammatory cytokines IL-6 and TNF-α, in both EOMG and LOMG. IL-6 promotes plasma cell differentiation, and autoantibody production as well as inhibiting the formation of Tregs^48^. Conversely, Tregs are known to suppress pro-inflammatory B cell responses and promote regulatory B cell differentiation^49^. In line with this, we found that IL-6^+^ B cell frequency correlated inversely with Treg frequencies, which were significantly reduced in refractory disease (but not the SIS cohort), in keeping with a previous study^50^. Importantly, Treg frequencies negatively correlated with disease severity and quality of life scores, highlighting their potential as a biomarker for disease activity. Of interest, anti-IL-6 treatment has shown benefit for MG in a recently published randomised controlled trial^51^, though Treg expansion strategies are also worthy of future study. Promoting immune tolerance confers lower risk than widespread immunosuppression, and can be done in a number of ways involving promoting regulatory innate immune cells^52,53^, possibly via vitamin D supplementation^54^, or adoptive transfer of Tregs which has shown therapeutic benefit in the mouse model of MG^55–57^. Therapies promoting antigen-specific tolerance in AChR-MG have also been investigated in early studies^58–63^. However, there are multiple disease-associated AChR-Ab epitopes^64^, which suggests that these strategies would need to be tailored to the individual.

In addition, we identify reduced frequency of DCs in refractory MG. A previous study showed decreased frequencies of pDCs, known to promote Treg differentiation^65^, in MG compared to controls, with a reduced pDC:mDC ratio^66^. Whilst a change in pDC:mDC ratio was not observed within our patient cohort, we identified a reduction in circulating pDCs and mDCs in refractory disease. pDC frequency also correlated with Treg frequencies, and inversely with disease severity and quality of life scores in MG patients. This is in contrast to a recent study reporting higher frequencies of circulating pDCs and mDCs in MG compared to control^19^. Notably, that study contained an overwhelming majority of patients with mild disease (MGFA class I cases), with low numbers of those with severe disease.

In agreement with previous findings, our data shows no differences in the frequency of naïve, memory, or effector T cells in patients with MG compared to controls, in relation to treatment or severity^67^. However, in contrast to other studies reporting an expansion of Tfh cells, we observe a nominal reduction in patients compared to controls^39^. The prior study defined Tfh as CD3^+^CD4^+^CXCR5^+^ cells, whereas we defined Tfh cells more robustly as CD3^+^CD4^+^CXCR5^+^CD45RA^-^PD-1^+^ cells. This subset comprises a small population of the circulating T cells, and changes seen in the circulation may not correspond to those in lymphoid organs.

Expansion of monocytes was seen in disease compared to controls, particularly in those with ongoing symptoms. Increased circulating classical monocyte frequencies may be promoting ongoing MHC-II antigen presentation, but also have roles in inflammatory cytokine secretion and promotion of B cell survival^68^. Both monocytes and DCs displayed reduced CD86 expression in those stable on immunosuppression, for monocytes this distinguished them from those who had not responded to immunosuppression (the refractory group). CD86 is the ligand for CD28 and CTLA-4 on T cells, and therefore reduced expression may result in impaired T cell activation and regulatory function^69^.

A previous study examining NK cell frequencies in MG showed elevated frequencies in those in remission compared to disease exacerbation and controls^70^. In agreement with this study, we report a trend to higher frequencies of NK cells in the SNIS cohort, suggesting they may play a protective role. Furthermore, other studies demonstrated a reduction in CD56^+^CD16^+^ NK cells, which correlated negatively with disease severity^14^, and an expansion of CD56dimCD16dim/- NK cells in MG patients with a high disease burden, due to prior ADCC activity^71^. However, we did not identify differences in NK cell subsets, perhaps due to differences in therapies used by participants.

Circulating levels of total C3, C5, and the inhibitory molecule clusterin were elevated in the refractory cohort, correlating positively with disease severity and quality of life scores, positioning them as additional potential biomarkers of disease activity. Although, other studies have reported varying results of complement levels, this might be contributed to by a number of confounding factors, including laboratory methodology and participant characteristics^22–25^. Elevated serum C5a levels have previously been found to correlate with clinical severity^25^. Conversely, another study found a negative correlation with C3 and disease severity, proposed to be due to increased complement consumption^22^. We did not see a difference in complement proteins between EOMG and LOMG, in contrast to a previous study that reported lower C3 and C4 levels in EOMG^23^. Given the known heterogeneity of complement activation by AChR-Abs^72^, prospective study of these complement markers prior to complement inhibitor therapies would allow assessment of their validity in predicting outcome for targeted treatment.

We demonstrated higher intensity of expression of complement receptors 1 (CD35) and 2 (CD21) on B cells, despite reduced frequencies of CD35^+^ B cells in disease, which may result in lower regulatory effects^30,31,73^. Notably, the expression of CD55 and CD46 on T cells was also increased in MG, with the greatest difference on CD8 T cells in refractory disease. CD55 and CD46 function to inhibit the complement cascade, but ligation also has functional effects on T cells^35^. CD55 ligation on T cells reduces IFN-γ and IL-2 production, and increases IL-10 production during antigen stimulation^74^, as well as having regulatory effects on CD8 T cell responses^75^. We found that inhibition of CD55 in CD4 T cells reduced TNF-α production, further supporting its role in promoting pro-inflammatory cytokine release. Whereas CD46 binding results initially in IFN-γ production by CD4 T cells^76^, subsequent expansion of effector cells results in IL-2 accumulation, which then promotes Treg development^77^. This switch to IL-10 production does not occur in CD8 T cells, and therefore the effects of CD46 on CD8 T cells are predominantly pro-inflammatory^78^. CD46 signalling has been shown to be dysregulated in several autoimmune conditions, with lack of Treg differentiation upon CD46 co-stimulation^77,79^. We found a disconnect between elevated CD46 expression and Treg frequencies, suggesting that similar CD46 dysregulation may also occur in MG, warranting further mechanistic investigation.

Our interpretation of the immune changes relevant to refractory disease were corroborated by unsupervised clustering within principal component analysis. The refractory-rich cluster displayed a distinct immune signature, characterised by several alterations including expanded memory B cells, enhanced CD55 expression on T cells, and low frequency of Tregs. Of note, all three non-refractory members of this cluster had EOMG, which could have, in-part, been relevant to some of the features seen.

Previous immune profiling relating to rituximab treatment for MG focuses on B cell repopulation, rather than baseline predictive markers. Early use of rituximab is previously reported as an indication of good response^80^, a finding we replicated here. We found that overall B cell frequency was <3% in all non-responders (those with a less than 50% reduction in disease severity score), and >3% in those with a ≥50% improvement. Although patient numbers are limited, this suggests that those early in disease will respond better than those with low B cell frequencies due to longstanding immunosuppression. Additionally, rituximab non-responders displayed the highest expression of CD55 on CD8 T cells, suggesting increased complement activity in these participants who may have benefitted from complement targeted therapy.

Following rituximab, the residual B cell population contained a high proportion of plasmablasts, likely because B cells lose CD20 expression as they differentiate into plasmablasts and plasma cells. This highlights the disadvantage of targeting CD20, as opposed to CD19, CD38, or B cell maturation antigen (BCMA). An anti-CD19 monoclonal antibody has recently demonstrated efficacy^81^, and trial results for plasma-cell targeting therapies are eagerly anticipated^82,83^.

The remaining non-depleted B cells were of an altered phenotype, with low BAFF-R, CD21 and CD35 expression. BAFF-R expression has been shown to be reduced on CD19 cells following rituximab previously^84^. Importantly, Tregs are known to expand following rituximab^50^, a finding corroborated in this study.

### Limitations of Study

The limitations of this study reflect those of many studies of MG. The wide heterogeneity of this rare disease makes it difficult to recruit to very specific cohorts, whilst trying to extrapolate the findings to a wider population. Here we attempted to limit this heterogeneity by only including those with AChR-ab-positive MG, with no thymoma or other autoimmune disease, however other potential confounding factors such as differences in age, sex, ethnicity, socio-economic status and other co-morbidities could have influenced the findings. Although power calculations were carried out to develop the protocol, recruitment fell short of planned numbers, and larger independent cohorts to validate these findings are required. The small numbers of refractory participants makes interpretation of differences between the responders and non-responders challenging. Additionally, as a cohort study, it remains unknown whether the changes seen within the refractory cohort have evolved over time or are present at disease onset to predict outcome. Prospective recruitment of those with severe disease prior to initiation of immunotherapy is challenging, nevertheless, these findings may constitute the basis for future studies, to validate these findings as predictive biomarkers.

### Summary

In summary, significant alterations to the innate and adaptive immune systems are observed in AChR-MG, particularly in refractory disease. Frequency of memory B cells or Tregs, or circulating C3, C5 or clusterin levels are potential prognostic markers that are worth further prospective evaluation. Our work suggests that future treatment strategies directed at plasma cells or at promoting immune tolerance mechanisms are likely to be beneficial in refractory disease.

## Acknowledgments

We would like to acknowledge the study participants for their contribution, the University of Manchester Flow Cytometry Core Facility, Holly Sedgwick, Maana Layeghi, Halima Ali-Shuwa, Rinal Sahputra, Bethany Potts, and Sara Kirkham for assistance and technical expertise and B. Paul Morgan and Tracy Hussell for intellectual input.

KD, PW, and LL performed statistical analyses, with oversight from MM. KD and PW performed the experiments. KD, MM, LL, and PW had unrestricted access to all data. KD and MM prepared the first draft of the manuscript, all authors read, edited, and approved the final draft and take responsibility for the content.

## Resource availability

Lead contacts: Further information and requests for resources and reagents should be directed to and will be fulfilled by the Lead Contacts, Madhvi Menon and Katherine Dodd

Materials availability: This study did not generate new unique reagents. Data and code availability:

◦ This study did not generate any original computer code.
◦ Data will be made available upon written proposal and data transfer agreement
◦ Any additional information required to reanalyze the data reported in this work paper is available from the Lead Contact upon request

## Author Contributions

Conceptualization: KCD, JSu, MM; Methodology: KCD, KB, JKLH, LL, MIL, JSp, SV, WMZ, JSu, PW, MM; Investigation: KCD, KB, JKLH, LL, MIL, JSp, SV, WMZ, JSu, PW, MM; Visualization and Data Analysis: KCD, LL, PW, MM; Funding acquisition: KCD, JSu; Project administration: KCD; Supervision: JSu, MM; Writing – original draft: KCD, MM; Writing – review & editing: KCD, KB, JKLH, LL, MIL, JSp, SV, WMZ, JSu, PW, MM

## STAR METHODS

### EXPERIMENTAL MODEL AND STUDY PARTICIPANT DETAILS

#### Ethics

This study was approved by Health Research Authority and Health Care Research Wales and NHS Research Ethics Committee (21/NW/0188). Written informed consent was obtained for all enrolled participants.

#### Study Design

Participants were recruited from the Northern Care Alliance NHS Foundation Trust, the Walton Centre NHS Foundation Trust, Newcastle Hospitals NHS foundation Trust, Oxford, University College London NHS Foundation Trust, and Imperial NHS foundation Trust. Case Record Forms and clinical assessment were completed by the neurology clinical research fellow with each participant at baseline and follow-up. All participants were able to give valid consent and were aged 18 to 80. The following inclusion criteria applied to each cohort.

Stable Immunosuppressed participants have a diagnosis of AChR positive myasthenia gravis (can be ocular, bulbar or generalised), with MGFA Post-intervention Status MM or better with no clinical relapse for 2 years, on either azathioprine or MMF along with ≤5mg/day of prednisolone. They had no prednisolone dose increase or decrease in past 12 months, and no increase in azathioprine or MMF dose for 2 years (allowing for cessation for up to 1 month). Other medications for other indications were allowed.

Stable Non-Immunosuppressed participants have a diagnosis of AChR positive myasthenia gravis (can be ocular, bulbar or generalised), with an MGFA Post-intervention Status MM or better on only low-dose cholinesterase inhibitors (≤120 mg pyridostigmine/day on average) for over two years. Other medications for other indications were allowed.

Refractory participants have a diagnosis of AChR positive myasthenia gravis (can be ocular, bulbar or generalised), and have been deemed eligible to be refractory to standard treatment and eligible for rituximab as per the NHS England criteria ^36^.

Treatment Naïve participants have a diagnosis of AChR positive myasthenia gravis, diagnosed in previous 6 months, and have not yet taken any form of immunosuppression.

Healthy Controls have no diagnosis of myasthenia gravis or other autoimmune condition that responds to standard immunosuppressive therapy (e.g. RA, inflammatory bowel disease).

Exclusion criteria for all cohorts included co-existing autoimmune condition for which azathioprine or mycophenolate mofetil are treatments (e.g. inflammatory bowel disease, RA, neuromyotonia), currently undergoing treatment for solid organ or haematological malignancy, or previous thymoma, or a clinical frailty scale ≥6.

58 participants were included. Demographics and clinical features of the included participants can be seen in supplementary table 1. Participants’ information on sex, age, and race was physician-reported. Association of results with sex, age and race was not performed throughout due to limited sample size. Information on gender and socioeconomic status was not collected and therefore any associations could not be analysed.

## METHOD DETAILS

### Sample Processing

Whole blood was collected in two EDTA tubes and processed within 12 hours of collection. Plasma was isolated from one tube via centrifugation of whole blood at 2000*g* for 10 minutes at room temperature. PBMCs were isolated from a separate tube using SepMate tubes (StemCell #85460), and Ficoll-Paque Plus (GE Healthcare # GE17-1440-03) via density gradient centrifugation (1200*g* for 10 minutes at room temp). Cells were washed twice in FACS buffer (PBS w/o Ca^2+^/Mg^2+^, 2%v/v FCS, 5 mM EDTA) prior to counting, using typtan blue and a countess II automated cell counter (Thermo Fisher Scientific). PBMCs were stored in FCS with 10% DMSO at -80°C for up to 2 weeks and then transferred to liquid nitrogen or -150°C storage.

### Flow Cytometry

Samples were thawed by warming at 37°C for 30 seconds then rapidly mixing with warmed complete media. Cells were washed in PBS then stained with Zombie UV™ (Biolegend, London UK) 1:500 for 15 minutes in PBS. Non-specific binding was blocked using Human FcR Blocking Reagent (Miltenyi, Bisley, UK) at 1:50 for 15 minutes. Extracellular antibodies were added at 1:50 for 30 minutes. Antibody information is provided in supplementary table 2. Cells were then fixed and permeabilised using FOXP3/Transcription Factor Staining Buffer Set (Thermo Fisher, Altrincham, UK), washed in perm buffer and stained with intracellular markers at 1:100 overnight. Cells were acquired on the LSRFortessa (BD Biosciences, Swindon, UK) and data analysed using FlowJo (v10.0; BD Biosciences). If under 50,000 lymphocytes were acquired, then results were excluded from analysis.

### Cell stimulation

Thawed cells were stimulated with either CD40L (R&D Systems, Abbingdon, UK) 0.5 µg/ml CpG-B (Cambridge Bioscience, Cambridge, UK) 1 µM, or R848 (Resiquimod, InvivoGen Toulouse, France) 1 µg/ml for 48 hours followed by 2 μl/ml of stimulation cocktail (Thermo Fisher) in the presence of Brefeldin A (Biolegend) in the last four hours, then washed and stained for flow cytometric analysis.

Thawed cells were also stimulated with R848 as above or placed into a plate which had been pre-coated for 1h at 37°C in PBS containing 1 µg/ml of each anti-CD3/anti-CD28 (clone OKT3, Biolegend (cat#317326)/ clone CD28.6, eBioscience (cat#16-0288-85) respectively) in isolation or with anti-CD21 (clone 1048, BD Biosciences (cat#552727)) 10 µg/ml, rCD35 (Bio-Techne (cat#5748-CD)) 20 µg/ml or anti-CD55 (Rabbit poly-clonal, AbCam (cat#ab231061)) 25 µg/ml for 48 hours followed by 2 μl/ml of stimulation cocktail (Thermo Fisher) in the presence of Brefeldin A (Biolegend) in the last four hours, then washed and stained for flow cytometric analysis.

### Complement Immunoassays

Fifteen complement analytes were selected based on previous research showing changes in serum levels of complement biomarkers in patients with MG and the accessibility of in house enzyme linked immunosorbent assays (ELISA) ^85–88^, The marker set was selected to interrogate the classical (C1q, C3, C4), alternative (Properdin, Ba, iC3b, FH, FHR125, FHR4, FI, sCR1), and terminal (C5, C9, clusterin, TCC (terminal complement complex)) pathways. Details of the antibodies, protein standards, and assay conditions can be found in supplementary table 3. Plasma samples stored at -80°C were thawed just before assaying, briefly vortexed, and then diluted in phosphate-buffered saline containing 0.1% Tween-20 (PBST, Sigma-Aldrich) and 0.2% bovine serum albumin (BSA) or non-fat milk (NFM; properdin assay only). Capture antibodies were fixed onto 96-well immunoplates (Thermo Fisher) overnight at 4°C, with concentrations ranging from 2–20 µg/ml in 50 µl/well of carbonate-bicarbonate buffer (pH 9.6). Wells were blocked with 2% BSA or NFM in PBST (100 µl/ well) and incubated for 1 h at 37°C, washed once with PBST, and plasma samples or protein standards diluted in buffer (0.2% BSA or NFM PBST (50 µl/ well) added in duplicate at a suitable dilution (supplementary table 3). Plates were incubated for 1h at RT or 37°C, washed three times and detection antibodies added at appropriate concentration (1–5 µg/ml) in 50 µl/well 0.2% BSA or NFM in PBST for 1h at RT or 37°C. In assays where the detection antibody was not directly labelled, an HRP-labelled secondary antibody (anti-mouse or anti-rabbit IgG, as applicable, Jackson ImmunoResearch, West Grove, USA) or for biotinylated antibodies, Streptavidin-HRP (R&D Systems) was used. Finally, plates were washed and developed using O-phenylenediamine dihydrochloride (OPD) substrate (Sigma-Aldrich) for 3–15 min, followed by adding 50 µl/well 5% H_2_SO_4_ to quench the reaction. Absorbances were read at 492 nm using Infinite F50 microplate reader (Tecan, Reading, UK). Intra- and inter-assay coefficient of variation were below 10% for all assays.

## QUANTIFICATION AND STATSTICAL ANALYSIS

### Sample Size

Sample size was calculated on previous unpublished data comparing CD19 count between those on effective immunosuppression and those not; for a power of 80% and a significant level of 5%, 23 participants were required in each group and we therefore attempted to recruit 24 to each cohort.

### Statistical Analysis

Statistical analysis was undertaken using GraphPad PRISM (version 10.0). All data underwent normality testing (Shapiro-Wilk test). Comparison between disease and control was with student’s t-test (parametric data) or Mann-Whitney (non-parametric). Comparison between cohorts was with one-way ANOVA with Tukey multiple comparison test (parametric) or Kruskal-Wallis with Dunn’s multiple comparison test (non-parametric). Results are presented as individual data points with medians. For correlation Pearson’s correlation (for parametric) or Spearman’s rank correlation coefficient test (for non-parametric data). In all cases, a p value of ≤ 0.05 was considered significant. Where no statistical difference is shown there was no significant difference. Details of statistical tests and definitions of n can be found in each figure legend. The bubble plot in figure 5 was created using R (version 4.5.2).

### Principal Component Analysis (PCA) and K-mean Clustering in PCA Space

Flow-cytometry markers were screened for missingness at the variable level (using Python version 3.13.5); any marker with >30% missing data was excluded (six markers removed: TNF-α⁺ B cells with R848, 74%; TNF-α⁺ B cells with CpG-B, 69%; IL-6⁺ B cells with CpG-B, 69%; IL-6⁺ B cells with R848, 67%; TNF-α⁺ B cells with CD40L, 64%; IL-6⁺ B cells with CD40L, 64%). Where the remaining analyses required complete data, remaining missing entries were then imputed using the per-marker median across samples, and all retained markers were standardised to z-scores (mean = 0, SD = 1).

Principal component analysis (PCA) was fitted to the z-scored matrix and the first two components (PC1, PC2) were used for visualisation. Unsupervised clustering was performed in the PC1–PC2 plane using K-means with a pre-specified number of clusters *k*; separation quality was summarised by the silhouette score. For each cluster, a confidence ellipse was drawn in PC space, centred at the cluster mean, oriented by the eigenvectors of the within-cluster covariance, and scaled to approximate a 95% contour.

For marker-level inference, a single target cluster from the K-means solution was compared with all other samples. For every marker, we computed Cohen’s 𝑑 (standardised mean difference with a pooled standard deviation) contrasting the target cluster against the remainder and additionally obtained two-sided Mann–Whitney U p-values. Markers were deemed different if they satisfied both criteria: 𝑝 < 0.05 and |𝑑| ≥ 0.5. To summarise multivariate patterns, we plotted a heatmap of mean z-scores for the target cluster versus others (rows) across markers (columns); where helpful, marker columns were hierarchically ordered to group similar response profiles.

## ADDITIONAL RESOURCES

This study was registered on clinicaltrials.gov ID: NHS001843.

