## Supplemental for "Lymphocyte alterations and elevated complement signalling are key features of refractory myasthenia gravis"

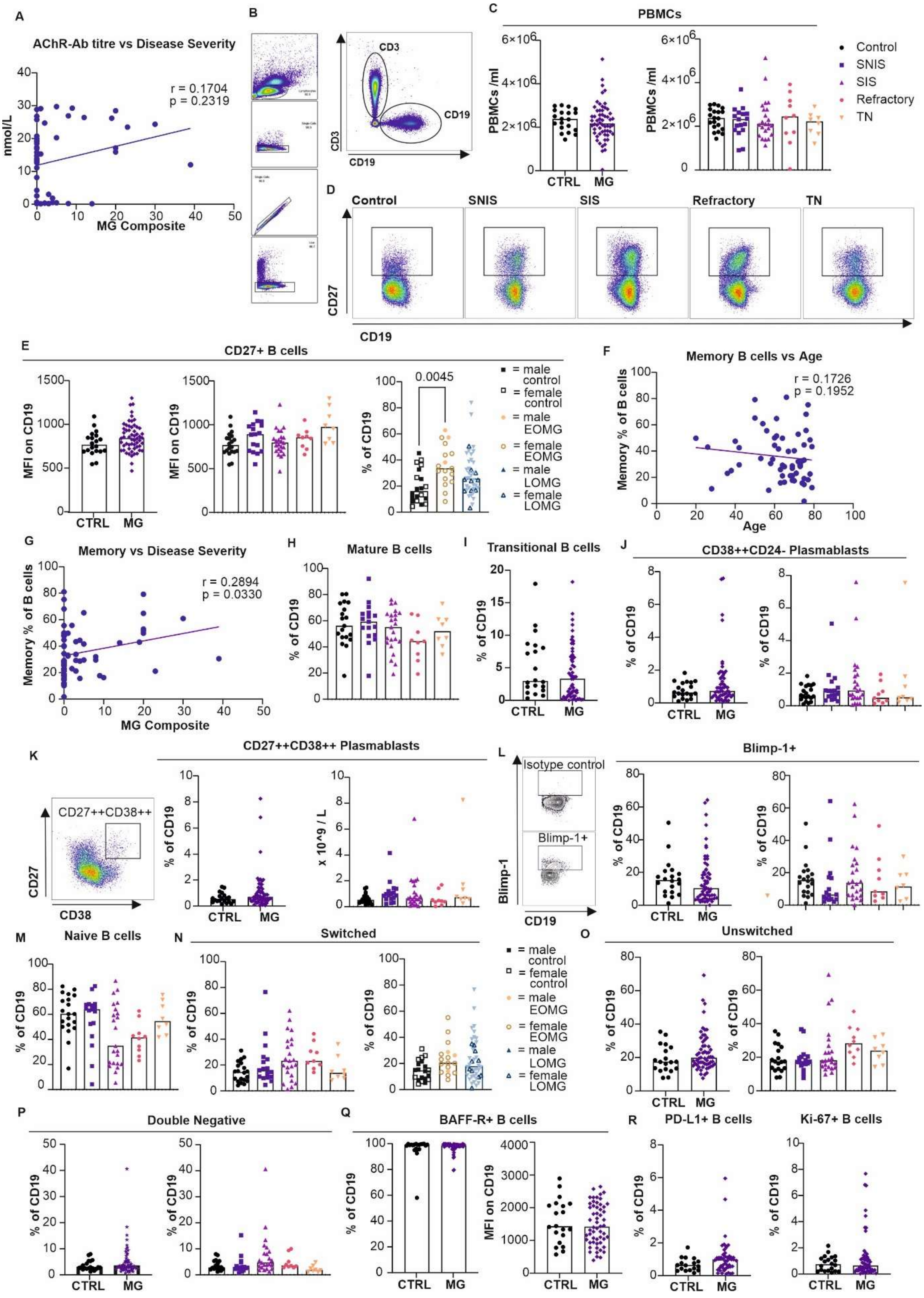


**Supplementary Figure 1.**

**Fig S1. Additional alterations to other B cell subset frequencies in AChR-MG, related to Figure 1.**

**(A)** AChR titre correlated with disease severity (n = 51). **(B)** Representative FACS gating strategy for CD19 live lymphocytes in PBMCs. **(C)** Graphs showing PBMC count /ml of blood in control (n = 20) compared to MG (n = 58), and between subgroups; control (n-20), SNIS (n = 17), SIS (n = 23), refractory (n = 10), and TN (n = 8). **(D)** Representative FACS plots for $CD27^+$ B cells for control, SNIS, SIS, refractory and TN cohorts. **(E)** Graphs showing CD27 MFI on B cells in control (n = 20) compared to MG (n = 57), between subgroups Control (n = 20), SNIS (n = 17), SIS (n = 22), refractory (n = 10), and TN (n = 8), and split into early- (n = 17) or late-onset MG (n = 40). **(F)** Graph showing simple linear regression between memory B cell frequency and age (n = 57). **(G)** Graphs showing simple linear regression between memory B cell frequency and disease severity (n = 57). **(H)** Graph showing mature B cell frequency within $CD19^+$ B cells in subgroups; control (n = 20), SNIS (n = 17), SIS (n = 22), refractory (n = 9), and TN (n = 8). **(I)** Graph showing transitional B cell frequency within $CD19^+$ B cells in Control (n = 20) compared to MG (n = 57). **(J)** Graphs showing $CD38^{++}CD24^-$ plasmablast frequencies in control (n = 20) compared to MG (n = 57), and between subgroups; control (n-20), SNIS (n = 17), SIS (n = 22), refractory (n = 9), and TN (n = 8). **(K)** FACS plot demonstrating $CD27^{++}CD38^{++}$ gating, gated on CD19+ B cells, and graphs showing $CD38^{++}CD27^{++}$ plasmablasts frequencies in control (n = 20) compared to MG (n = 57), and between subgroups; control (n = 20), SNIS (n = 17), SIS (n = 22), refractory (n = 9), and TN (n = 8). **(L)** FACS plots demonstrating Blimp-1 gating strategy, gated on $CD19^+$ B cells, and graphs showing Blimp-1+ B cell frequency in control (n = 20) compared to MG (n = 57), and between subgroups; control (n-20), SNIS (n = 17), SIS (n = 22), refractory (n = 10), and TN (n = 8). **(M)** Graphs showing naïve ($IgD^+CD27^-$) B cells in subgroups; control (n = 20), SNIS (n = 17), SIS (n = 22), refractory (n = 9), and TN (n = 8). **(N)** Graph showing switched B cell frequencies in subgroups; control (n = 20), SNIS (n = 17), SIS (n = 22), refractory (n = 9), and TN (n = 8), and when split into early- (n = 17) or late-onset MG (n = 40). **(O)** Graphs showing unswitched memory and **(P)** double negative (IgD-CD27-) B cells in control (n = 20) compared to MG (n = 57), and between subgroups; control (n = 20) SNIS (n = 17), SIS (n = 22), refractory (n = 9), and TN (n = 8). **(Q)** Graphs showing frequencies and expression of $BAFF\text{-}R^+$ B cells, in control (n = 20) compared to MG (n = 57). **(R)** Graphs showing frequencies of $PD\text{-}L1^+$ and $Ki\text{-}67^+$ B cells, in control (n = 20) compared to MG (n = 57).

Graphs show individual participant data, with bar representing median values.

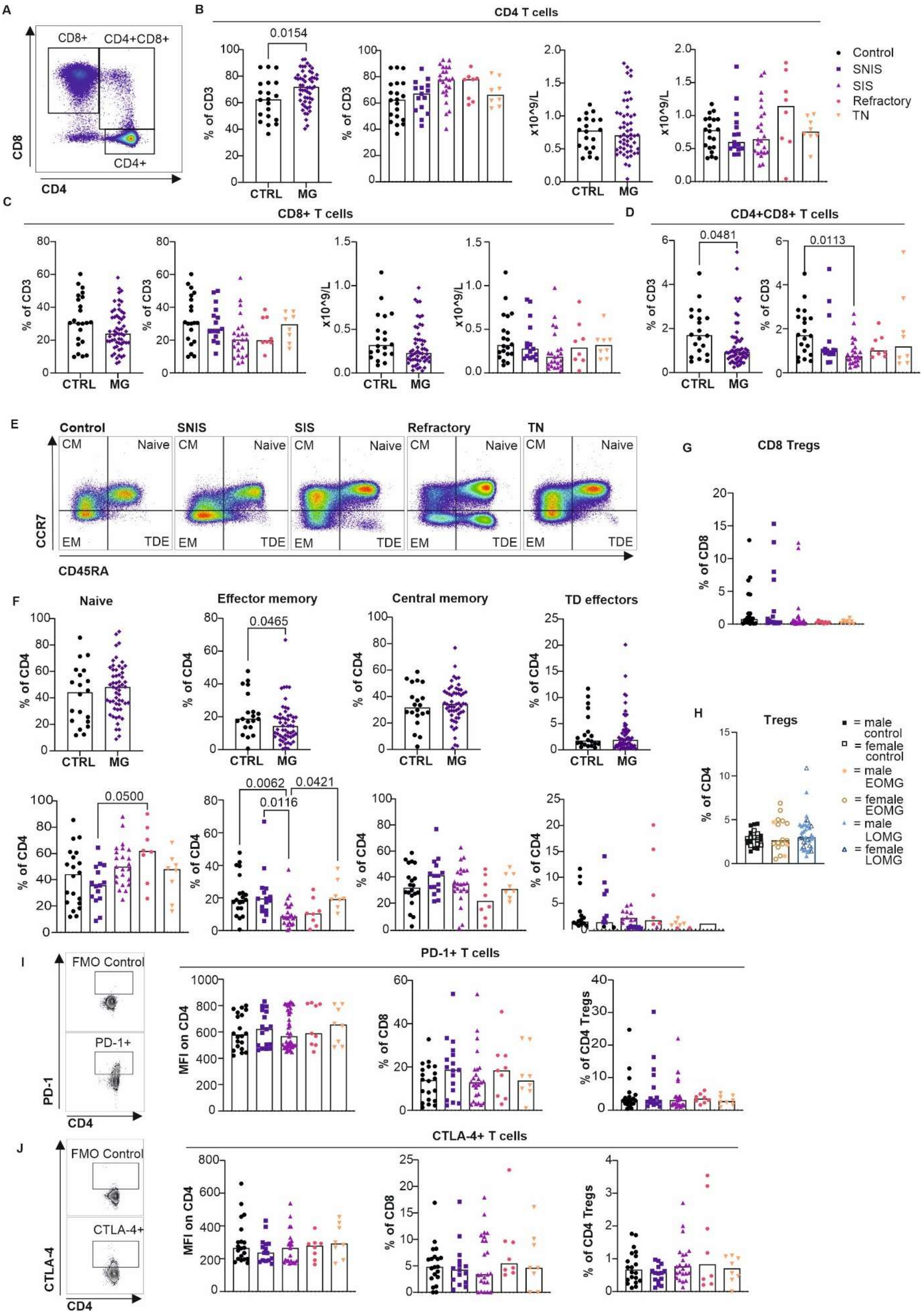


**Supplementary Figure 2.**

**Fig. S2. Alterations to T cell subsets in AChR-MG, related to Figure 3**.

**(A)** FACS plot showing $CD4^+$, $CD8^+$ and $CD4^+CD8^+$ gating, gated on CD3+ live lymphocytes in PBMCs. **(B)** Graphs showing CD4+ frequency and count in control (n = 20) compared to MG (n = 54), and between MG subgroups; SNIS (n = 15), SIS (n = 23), refractory (n = 8), and TN (n = 8). **(C)** Graphs showing CD8+ frequency and count in control (n = 20) compared to MG (n = 54), and between MG subgroups; SNIS (n = 15), SIS (n = 23), refractory (n = 8), and TN (n = 8). **(D)** Graphs showing CD4+CD8+ frequency in control (n = 20) compared to MG (n = 54), and between MG subgroups; SNIS (n = 15), SIS (n = 23), refractory (n = 8), and TN (n = 8). **(E)** Representative FACS plots demonstrating gating of naïve, central and effector memory and TD T cells. **(F)** Graphs showing naïve, effector and central memory and TD effector T cell frequency in control (n = 20) compared to MG (n = 54), as well as in subgroups; control (n = 20), SNIS (n = 15), SIS (n = 23), refractory (n = 8), and TN (n = 8). **(G)** Graph showing simple linear regression for Treg frequency against disease severity (n = 54) **(H)** Graph showing Treg frequencies in control (n = 20), compared to EOMG (n = 16) and LOMG (n = 33). **(I)** Graph showing frequency of Tfh cells in control (n = 20) compared to MG (n = 54). **(J)** FACS plot demonstrating gating strategy for $PD\text{-}1^+$ CD4 T cells, and graphs showing PD-1+ expression on CD4, and PD-1+CD4 Treg and CD8 T cell frequency in control (n = 20), SNIS (n = 15), SIS (n = 23), refractory (n = 8), and TN (n = 8). **(K)** FACS plot demonstrating gating strategy for $CTLA\text{-}4^+$ on CD4 T cells, and graphs showing $CTLA\text{-}4^+$ expression on CD4 T cells and $CTLA\text{-}4^+$ CD4 Treg and CD8 T cell frequency in control (n = 20), SNIS (n = 15), SIS (n = 23), refractory (n = 8), and TN (n = 8).

Graphs show individual participant data, with bar representing median values.

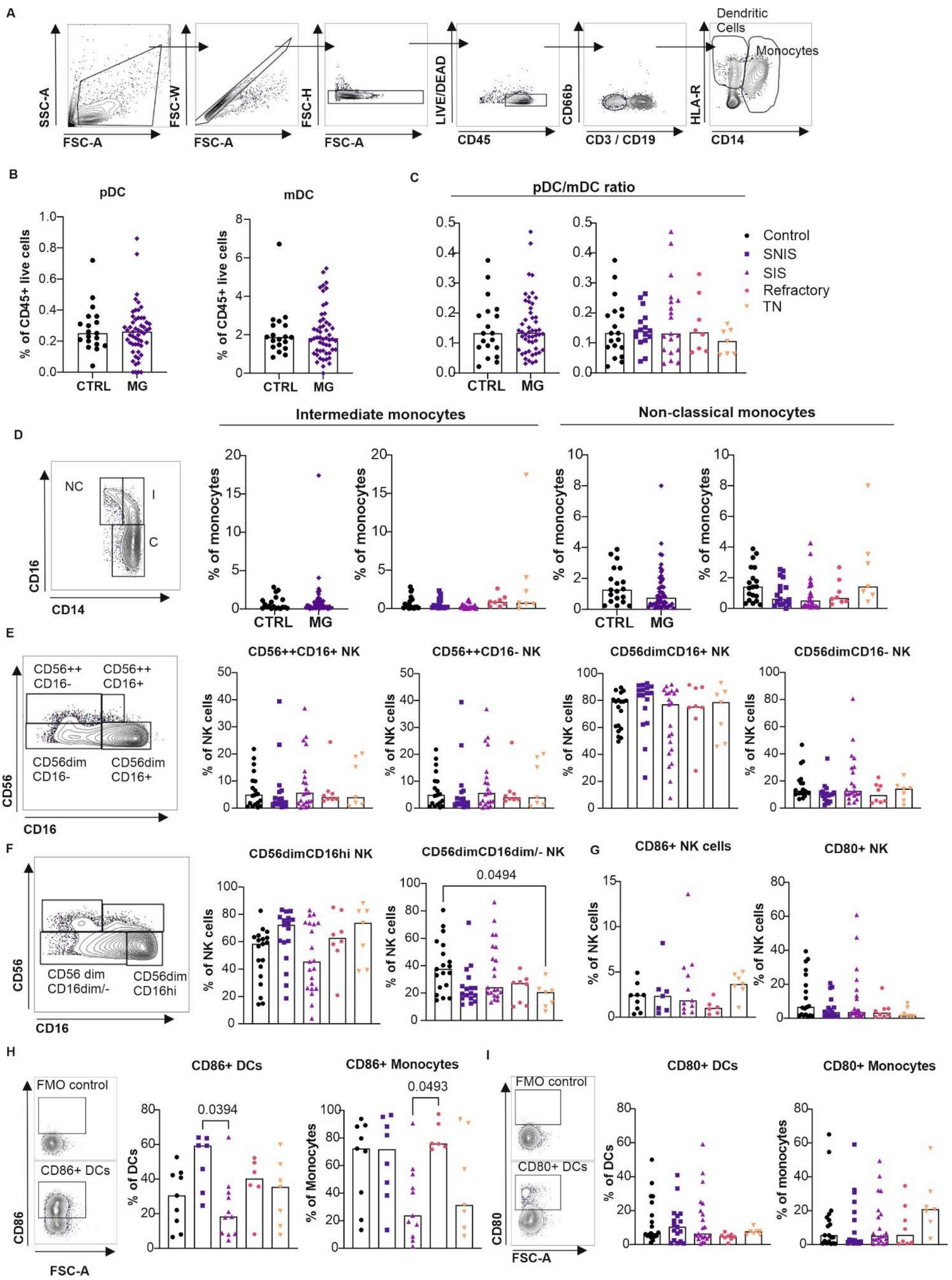


**Supplementary Figure 3.**

**Fig. S3. Alterations to myeloid cells in AChR-MG, related to Figure 4.**

**(A)** Representative FACS plots demonstrating gating strategy for dendritic cells and monocytes. **(B)** Graphs showing pDC and mDC frequency in control (n = 18) compared to MG (n = 48 ). **(C)** Graph showing DC frequency and pDC/mDC ratio in control (n = 18) compared to MG (n = 48 ), and pDC/mDC ratio in subgroups; control (n = 18), SNIS (n = 17), SIS (n = 19), refractory (n = 6), TN (n = 6). **(D)** Representative FACS plot demonstrating gating strategy for classical (C), intermediate (I) and non-classical (NC) monocytes and graphs showing frequencies of intermediate and non-classical monocytes in control (n = 18), compared to MG (n = 48), and in subgroups control (n = 18), SNIS (n = 17), SIS (n = 19), refractory (n = 6), and TN (n = 6). **(E)** Representative FACS plot demonstrating gating strategy for $CD56^{bright}CD16^{+}$, $CD56^{bright}CD16^{-}$, $CD56^{dim}CD16^{+}$ and $CD56^{dim}CD16^{-}$ NK cells, gated on $CD3^{-}CD19^{-}CD56^{+}$ live lymphocytes, and graphs showing frequencies of these populations for subgroups; control (n = 18), SNIS (n = 17), SIS (n = 19), refractory (n = 6), TN (n = 6). **(F)** Representative FACS plot demonstrating gating strategy for $CD56^{dim}CD16^{hi}$ and $CD56^{dim}CD16^{+/-}$ NK cells, gated on $CD3^{-}CD19^{-}CD56^{+}$ live lymphocytes, and graphs showing frequencies of these populations for subgroups; control (n = 18), SNIS (n = 17), SIS (n = 19), refractory (n = 6), TN (n = 6). **(G)** Frequencies of $CD86^{+}$ and $CD80^{+}$ NK cells in subgroups; control (n = 18), SNIS (n = 17), SIS (n = 19), refractory (n = 6), and TN (n = 6). **(H)** FACS plot demonstrating gating strategy using an FMO for CD86 on DCs, and graphs showing $CD86^{+}$ frequency in DCs and monocytes for subgroups; control (n = 18), SNIS (n = 17), SIS (n = 19), refractory (n = 6), and TN (n = 6). **(I)** FACS plots demonstrating gating strategy using an FMO for CD80 on DCs and graphs showing $CD80^{+}$ frequency in DCs and monocytes for subgroups; control (n = 18), SNIS (n = 17), SIS (n = 19), refractory (n = 6), and TN (n = 6).

Graphs show individual participant data, with bar representing median values.

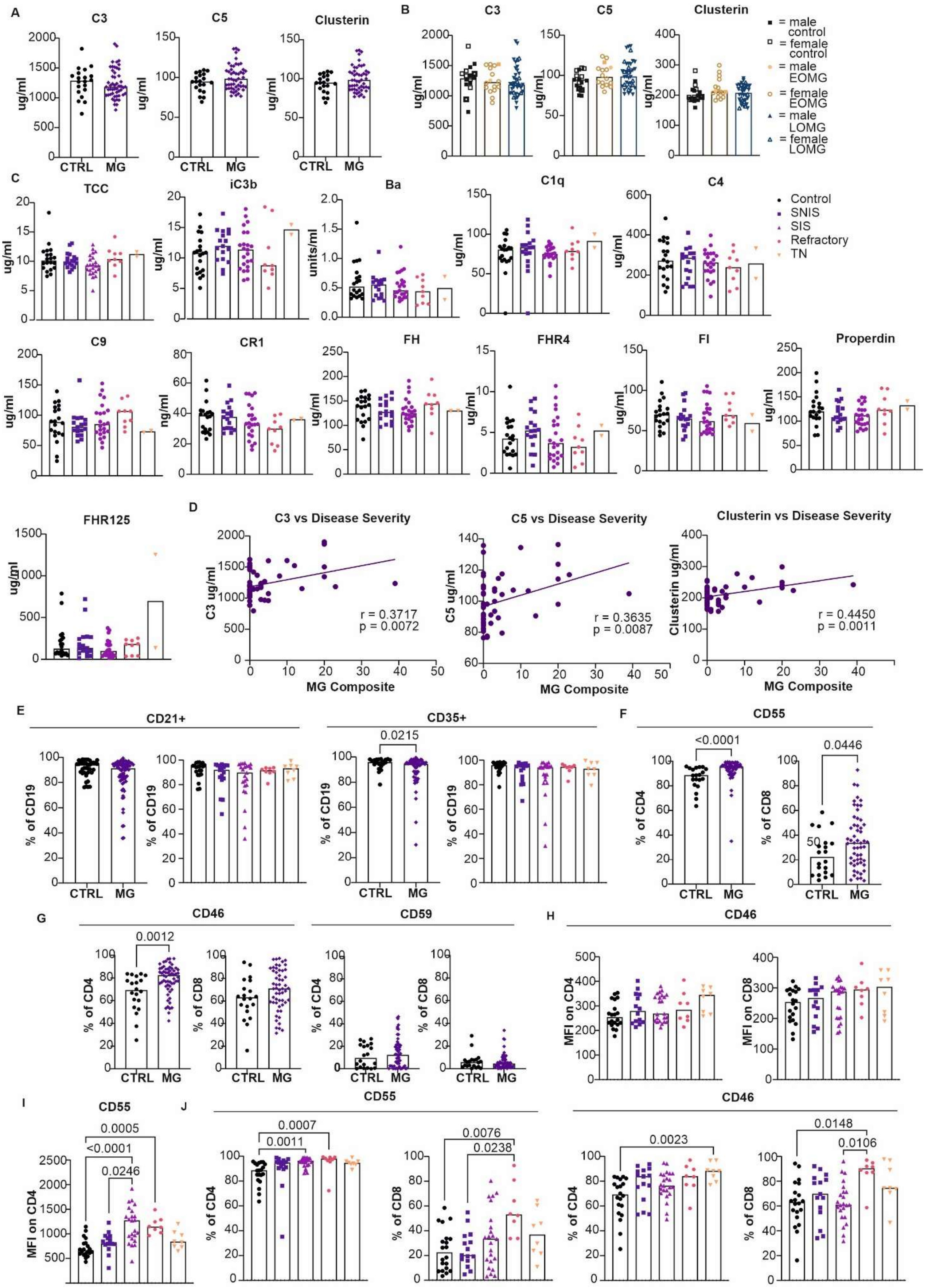

**Supplementary Figure 4.**

**Fig. S4. Complement proteins and lymphocyte complement receptors in AChR-MG, related to Figure 5.**

**(A)** Graphs showing frequencies of C3, C5, and clusterin in control (n = 20) compared to MG (n = 52). **(B)** Graphs showing C3, C5 and clusterin levels in control (n = 20), compared to EOMG (n = 16) and LOMG (n = 38). **(C)** Graphs showing frequencies of TCC, iC3b, Ba, C1q, C4, C9, CR1, FH, FHR4, FI, Properdin and FHR125 in control (n = 20) compared to MG (n = 52), and between subgroups control (n = 20), SNIS (n = 18), SIS (n = 23), refractory (n = 9), and TN (n = 2). **(D)** Graphs showing simple linear regression between C3, C5 and clusterin with disease severity (as measured by the MG Composite score) (n = 56). **(E)** Graphs showing frequency of $CD21^+$ and $CD35^+$ B cells in control (n = 20) compared to MG (n = 56) and in subgroups control (n = 20), SNIS (n = 17), SIS (n = 22), refractory (n = 9), and TN (n = 8). **(F)** Graphs showing $CD55^+$ CD4 and CD8 T cell frequency in control (n = 20) and MG (n = 54). **(G)** Graphs showing $CD46^+$ and $CD59^+$ CD4 and CD8 T cells in control (n = 20) compared to MG (n = 54). **(H)** Graphs showing CD46 expression on CD4 and CD8 T cells in subgroups control (n = 20), SNIS (n = 15), SIS (n = 23), refractory (n = 8), and TN (n = 8). **(I) (H)** Graph showing CD55 expression on CD4 T cells in subgroups control (n = 20), SNIS (n = 15), SIS (n = 23), refractory (n = 8), and TN (n = 8). **(J)** Graphs showing frequency of $CD55^+$ and $CD46^+$ CD4 and CD8 T cells in subgroups control (n = 20), SNIS (n = 15), SIS (n = 23), refractory (n = 8), and TN (n = 8).

Graphs show individual participant data, with bar representing median values.

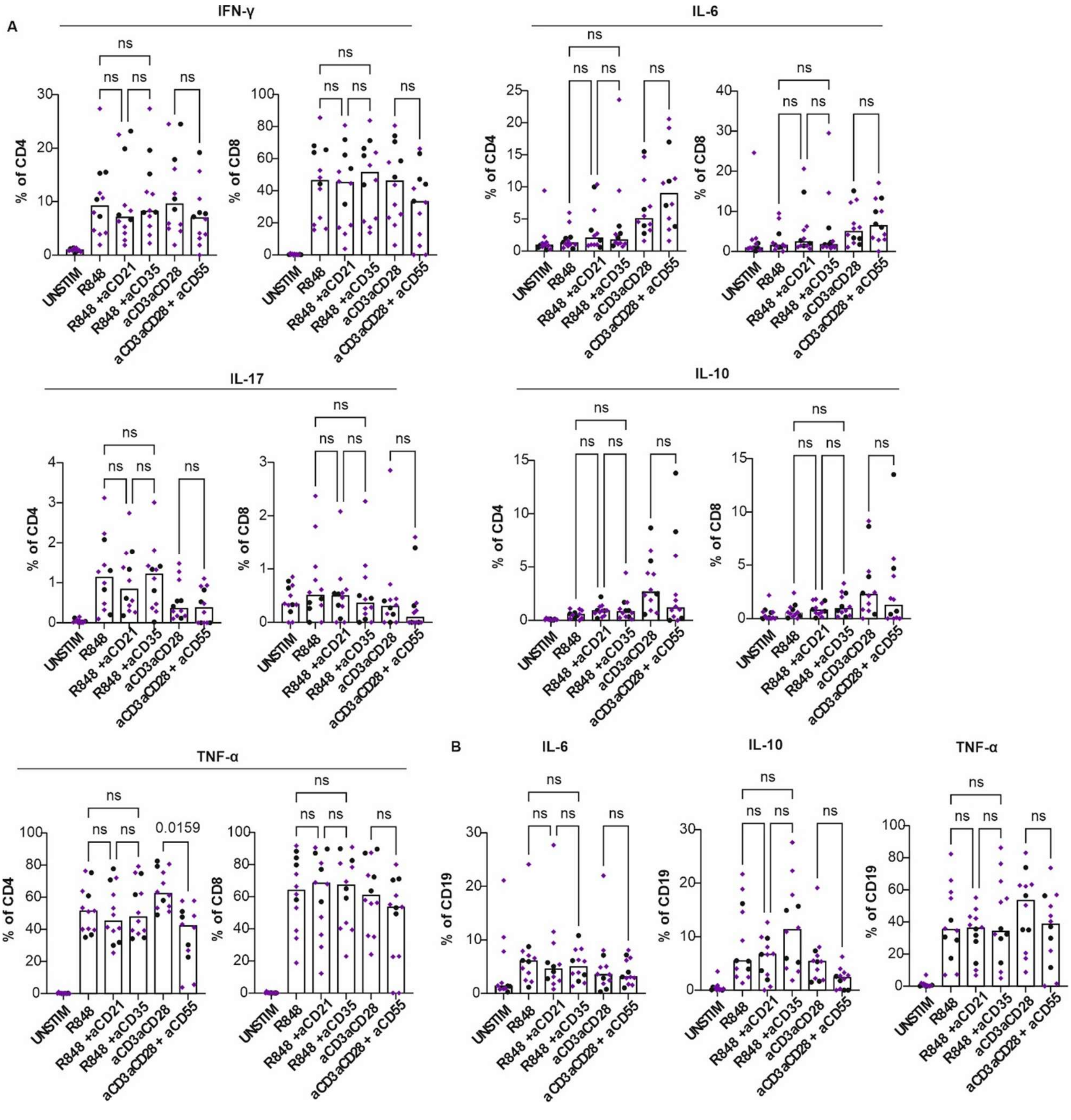


**Supplementary Figure 5.**

**Fig. S5. Lymphocyte cytokine production in the context of CD35, CD21 and CD55 inhibition, related to Figure 5.**

**(A)** Graphs showing IFN-γ, IL-6, IL-17, IL-10, TNF-α positive CD4 and CD8 T lymphocytes when unstimulated, when stimulated by R848 with or without anti-CD21 or anti-CD35, and when stimulated by aCD3aCD28 with or without anti-CD55. **(B)** Graphs showing IL-6, IL-10, and TNF-α positive CD19 lymphocytes when unstimulated, when stimulated by R848 with or without anti-CD21 or anti-CD35, and when stimulated by aCD3aCD28 with or without anti-CD55. Graphs show individual participant data, with bar representing median values. Comparisons are performed by one-way ANOVA (parametric data) or Friedman test (non-parametric).

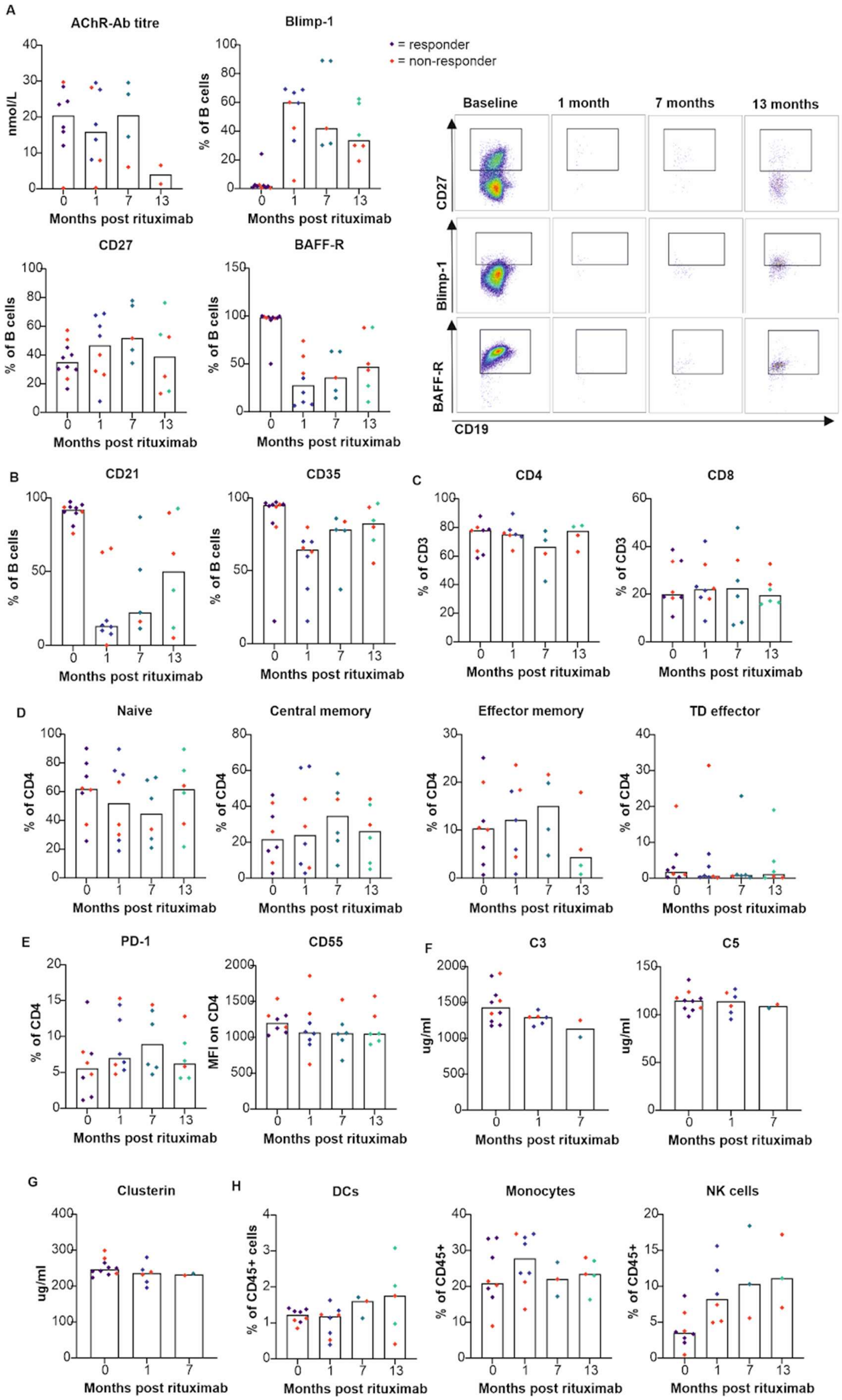


**Supplementary Figure 6.**

**Fig. S6. Prospective changes seen following rituximab therapy for refractory AChR-MG, related to Figure 7.**

**(A)** Graphs showing AChR tire and CD27$^+$, Blimp-1$^+$ and BAFF-R$^+$ B cell frequency in the refractory group at baseline (n = 8 AChR, n = 10 B cells) and 1 month (n = 8), 7 months (n = 4 AChR, n = 6 B cells), and 13 months (n = 2 AChR, n = 6 B cells) post rituximab, and representative FACS plots showing changes in CD27$^+$, Blimp-1$^+$ and BAFF-R$^+$ B cell frequency. **(B)** Graphs showing CD21$^+$ and CD35$^+$ B cell frequency in the refractory group at baseline (n = 10) and 1 month (n = 8), 7 months (n = 5), and 13 months (n = 6) post rituximab. **(C)** Graphs showing CD4 and CD8 T cell frequency at baseline (n = 10), to 1 month (n = 8), 7 months (n = 5), and 13 months (n = 6) post rituximab. **(D)** Graphs showing naïve, central memory, effector memory and TD effector T cell frequency at baseline (n = 10) and 1 month (n = 8), 7 months (n = 5), and 13 months (n = 6) post rituximab. **(E)** Graphs showing PD-1$^+$ CD4 T cell frequency and CD55 expression on CD4 T cells at baseline (n = 10) and 1 month (n = 8), 7 months (n = 5), and 13 months (n = 6) post rituximab. **(F)** Graphs showing C3 and C5 levels at baseline (n = 10) and 1 month (n = 6), and 7 months (n = 2) post rituximab. **(G)** Graph showing changes to clusterin levels at baseline (n = 10) and 1 month (n = 6), and 7 months (n = 2) post rituximab. **(H)** Graphs showing changes in monocyte, DC and NK cell frequency in the refractory group at baseline (n = 10), to 1 month (n = 8), 7 months (n = 4), and 13 months (n = 5) post rituximab.

Graphs show individual participant data, with bar representing median values. Red points denote "non-responders" to rituximab.

***Supplementary Table 1. Demographics and clinical characteristics of participants, related to STAR methods.***

| | ***Control*** | ***MG total*** | | | | |
|---|---|---|---|---|---|---|
| | | | **SNIS** | **SIS** | **Refractory** | **TN** |
| ***N*** | **20** | **58** | 17 | 23 | 10 | 8 |
| ***Age*** *(mean, years)* | **61.5** | **63.0** | 65.1 | 66.7 | 54.5 | 59.8 |
| ***Sex*** *(% female)* | **45%** | **41%** | 47% | 30% | 50% | 50% |
| ***BMI*** | **30.0** | **29.0** | 27.6 | 28.9 | 32.7 | 27.6 |
| ***Ethnicity*** | | | | | | |
| *Asian* | **10%** | **6.9%** | 5.9% | 4.3% | 10% | 12.5% |
| *Afrocaribbean* | **5%** | **3.4%** | 0% | 4.3% | 10% | 0% |
| *Caucasian* | **85%** | **89.7%** | 94.1% | 91.3% | 80% | 87.5% |
| ***Disease severity*** *(mean, MGC / 50)* | **N/A** | **5.2** | 1.1 | 1.2 | 19.7 | 7.25 |
| ***Disease-related quality of life*** *(mean, MG-QoL 15r /30)* | **N/A** | **7.6** | 1.1 | 4 | 22.3 | 13.6 |
| ***Disease duration*** *(mean, years)* | **N/A** | **9.1** | 11.0 | 12.3 | 5.3 | 0.36 |
| ***Prednisolone dose*** *(mean over past 3 months, mg/day)* | **0** | **4.4** | 0 | 0.8 | 23.8 | 0 |
| ***Other Immunosuppression*** | **Nil** | **26% AZA<br>22% MMF<br>1.7% MTX** | Nil | 57% AZA<br>43% MMF | 20% AZA<br>30% MMF<br>10% MTX | Nil |
| ***Co-morbidities*** | | | | | | |
| *T2DM* | **10%** | **26%** | 18% | 30% | 50% | 0% |
| *HTN* | **35%** | **41%** | 24% | 48% | 50% | 50% |
| *OA* | **20%** | **9%** | 6% | 0% | 20% | 25% |
| *Hight cholesterol* | **50%** | **17%** | 35% | 11% | 30% | 25% |
| *COPD / Asthma* | **30%** | **19%** | 18% | 17% | 20% | 25% |
| *Hypothyroidism* | **5%** | **5%** | 12% | 0% | 10% | 0% |
| ***AChR-Ab titre*** *(median, nmol/L, normal <0.45)* | **0.23** | **14.76** | 13.16 | 13.83 | 23.92 | 17.16 |
| ***Total lymphocyte count*** *(median, x10^9/L, normal range 1.0-4.0)* | **1.5** | **1.4** | 1.4 | 1.1 | 2.0 | 1.7 |

*Abbreviations: AChR-Ab, acetylcholine receptor antibody; AZA, azathioprine; BMI, body mass index; MTX, methotrexate; MG, myasthenia gravis; MGC, MG Composite; MMF, mycophenolate mofetil; SIS, stable immunosuppressed; SNIS, stable non-immunosuppressed; TN, treatment naïve.*

*Supplementary Table 2. Antibodies used in flow cytometry staining, , related to STAR methods*

| Fluorophore | Marker | Clone | Supplier | Catalog Number |
|---|---|---|---|---|
| **B cell panel** | | | | |
| AF488 | Blimp-1 | 646702 | R&D Systems | IC36081G |
| PerCP/Cy5.5 | CD38 | HIT-2 | Biolegend | 303522 |
| APC | CD21 | Bu32 | Biolegend | 254906 |
| AF700 | Ki67 | Ki-67 | Biolegend | 350530 |
| APC/Fire 750 | CD24 | ML5 | Biolegend | 311140 |
| BV421 | IgG | M1310G05 | Biolegend | 410704 |
| BV510 | IgM | MHM-88 | Biolegend | 314522 |
| BV605 | IgD | IA6-2 | Biolegend | 348232 |
| BV650 | CD3 | UCHT1 | Biolegend | 300468 |
| BV711 | CD274 (PD-L1) | 29E.2A3 | Biolegend | 329722 |
| BV785 | CD19 | HIB19 | Biolegend | 302240 |
| PE. | CD35 | E11 | Biolegend | 333406 |
| PE Dazzle 594 | CD27 | M-T271 | Biolegend | 329732 |
| PeCy7 | BAFF-R | 11C1 | Biolegend | 316920 |
| UV – 450 | Live / dead | Zombie UV Fixable Viability Kit | Biolegend | 423107 |
| **T cell panel** | | | | |
| FITC | FoxP3 | 206D | Biolegend | 320106 |
| PerCP/Cy5.5 | CD4 | OKT4 | Biolegend | 317428 |
| APC | CD59 | p282 | Biolegend | 304712 |
| AF700 | CXCR5 | J252D4 | Biolegend | 356916 |
| APC/Fire 750 | CD56 | 5.1H11 | Biolegend | 362554 |
| BV421 | CCR7 (CD197) | G043H7 | Biolegend | 353208 |
| BV510 | CD279 (PD-1) | EH12.2H7 | Biolegend | 329932 |
| BV605 | CD45RA | HI100 | Biolegend | 304134 |
| BV650 | CD3 | UCHT1 | Biolegend | 300468 |
| BV711 | CD25 | BC96 | Biolegend | 302636 |
| BV785 | CD8 | SK1 | Biolegend | 344740 |
| PE | CD55 (DAF) | JS11 | Biolegend | 311308 |
| PE Dazzle 594 | CTLA-4 (CD152) | L3D10 | Biolegend | 349922 |
| PeCy7 | CD46 (MCP) | TRA-2-10 | Biolegend | 352408 |
| UV – 450 | Live / dead | Zombie UV Fixable Viability Kit | Biolegend | 423107 |
| **Myeloid Panel** | | | | |
| FITC | CD14 | M5E2 | Biolegend | 301804 |
| PerCP/Cy5.5 | HLA-DR | L243 | Biolegend | 307630 |
| APC | CD123 | 6H6 | Biolegend | 306012 |
| AF700 | Ki67 | Ki-67 | Biolegend | 350530 |
| APC/Fire 750 | CD56 | 5.1H11 | Biolegend | 362554 |
| BV421 | CD66b | 6/40c | Biolegend | 392916 |
| BV510 | CD45 | HI30 | Biolegend | 304036 |

| | | | | |
|---|---|---|---|---|
| BV605 | CD16 | 3G8 | Biolegend | 302040 |
| BV 650 | CD19 | HIB19 | Biolegend | 302238 |
| BV650 | CD3 | UCHT1 | Biolegend | 300468 |
| BV711* | CD274 (PD-L1) | 29E.2A3 | Biolegend | 329722 |
| BV711* | CD86 | 29E.2A3 | Biolegend | 329722 |
| BV785 | CD11c | 3.9 | Biolegend | 301644 |
| PE | CD64 | 10.1 | Biolegend | 305008 |
| PE Dazzle 594 | CD303 (BDCA-2) | 201A | Biolegend | 354226 |
| PeCy7 | CD80 | 2D10 | Biolegend | 305218 |
| UV – 450 | Live / dead | Zombie UV Fixable Viability Kit | Biolegend | 423107 |
| **Stimulation Panel** | | | | |
| FITC | CD14 | M5E2 | Biolegend | 301804 |
| PerCP/Cy5.5 | HLA-DR | L243 | Biolegend | 307630 |
| APC | IL-10 | JES3-19F1 | Biolegend | 506807 |
| AF700 | CD4 | SK3 | Biolegend | 344622 |
| BV605 | CD3 | 3G8 | Biolegend | 302040 |
| BV 650 | CD19 | HIB19 | Biolegend | 302238 |
| PE | IL-6 | MQ2-13A5 | Biolegend | 501107 |
| Pe Cy7 | TNF-α | MAb11 | Biolegend | 502930 |
| UV – 450 | Live / dead | Zombie UV Fixable Viability Kit | Biolegend | 423107 |
| **Complement Receptor Stimulation Panel** | | | | |
| FITC | CD35 | E11 | Biolegend | 333404 |
| PerCP/Cy5.5 | CD4 | OKT4 | Biolegend | 317428 |
| APC | CD21 | Bu32 | Biolegend | 254906 |
| AF700 | CD38 | HIT2 | Biolegend | 303524 |
| APC/Fire 750 | CD24 | ML5 | Biolegend | 311140 |
| BV421 | IgG | M1310G05 | Biolegend | 410704 |
| BV510 | IgM | MHM-88 | Biolegend | 314522 |
| BV605 | IgD | IA6-2 | Biolegend | 348232 |
| BV650 | CD3 | UCHT1 | Biolegend | 300468 |
| BV711 | CD19 | HIB19 | Biolegend | 302246 |
| BV785 | CD8 | SK1 | Biolegend | 344740 |
| PE | CD55 (DAF) | JS11 | Biolegend | 311308 |
| PE Dazzle 594 | CD27 | M-T271 | Biolegend | 329732 |
| PeCy7 | CD46 (MCP) | TRA-2-10 | Biolegend | 352408 |
| UV – 450 | Live / dead | Zombie UV Fixable Viability Kit | Biolegend | 423107 |
| **Complement Receptor Inhibitor / Stimulation Panel** | | | | |
| PerCP/Cy5.5 | CD4 | OKT4 | Biolegend | 317428 |
| APC | CD3 | BW/264/56 | Miltenyi | 130-113-125 |
| AF700 | IL-17A | BL168 | Biolegend | 512318 |
| BV421 | IgG | M1310G05 | Biolegend | 410704 |
| BV510 | IgM | MHM-88 | Biolegend | 314522 |
| BV605 | IgD | IA6-2 | Biolegend | 348232 |

| | | | | |
|---|---|---|---|---|
| BV650 | CD19 | HIB19 | Biolegend | 302238 |
| BV711 | IFN-G | 4S:B3 | Biolegend | 502540 |
| BV785 | CD8 | SK1 | Biolegend | 344740 |
| cellPE | IL-6 | MQ2-13A5 | Biolegend | 501107 |
| PE Dazzle 594 | IL-10 | JES3-19F1 | Biolegend | 506812 |
| PeCy7 | TNF-α | MAb11 | Biolegend | 502930 |
| UV - 450 | Live / dead | Zombie UV Fixable Viability Kit | Biolegend | 423107 |

*****interchanged after 2 batches

*Supplementary Table 3. Antibodies, standards and sample dilutions used in complement ELISA, , related to STAR methods*

| Assay | Capture antibody | Detection antibody | Protein | Detection range (ng/ml) | Dilution factor |
|---|---|---|---|---|---|
| Ba | mAb D22/3 (Hycult) | mAb P21/15-HRP (Hycult) | Ba (Comptech) | 8.5–550 | 1:4 |
| iC3b | mAb clone 9 | mAb bH6 -HRP (Hycult) | iC3b (Comptech) | 23–1500 | 1:40 |
| TCC | mAb aE11 (Hycult) | mAb E2 anti-C8-biotin | TCC | 39–2500 | 1:40 |
| C1q | mAb 9H10 | Rabbit anti-C1q | C1q | 23–1500 | 1:16000 |
| C3 | mAb 2898 (Hycult) | mAb clone 3-HRP (Hycult) | C3 | 31–2000 | 1:2000 |
| C4 | Rabbit anti-C4 | Rabbit anti-C4-HRP | C4 (Comptech) | 15–1000 | 1:4000 |
| C5 | mAb 10B6 | mAb 4G2-HRP | C5 | 39–2500 | 1:500 |
| C9 | mAb B7 | Rabbit anti-C9-biotin | C9 | 3–200 | 1:4000 |
| Clusterin | mAb 2D5 | mAb 4C7-HRP | Clusterin | 31–2000 | 1:1000 |
| sCR1 | Rabbit anti-sCR1 | mAb MBI35-HRP | sCR1 | 0.4–25 | 1:2 |
| FH | mAb OX24 | mAb 35H9-HRP | FH | 15–1000 | 1:4000 |
| FHR4 | mAb 4E9 | mAb clone 150-HRP | FHR4 | 15–1000 | 1:100 |
| FHR125 | mAb MBI125 | clone 35H9-HRP | FHR125 | 156–10,000 | 1:100 |
| FI | mAb 7B5 | Rabbit anti-FI | FI | 15–1000 | 1:500 |
| Properdin | mAb 9.3.4 (Hycult) | mAb clone 2.9 (Hycult) | Properdin (Comptech) | 6–400 | 1:1600 |

*All antibodies were prepared in house unless stated otherwise specified for each analyte. mAb; monoclonal antibody, HRP, horseradish peroxidase, Comptech; Complement Technology Inc.,TCC, terminal complement complex; FH, factor H; FHR125, FH-related proteins 1, 2, and 5; FHR4, FH-related protein 4; sCR1, soluble complement receptor type 1; FI, factor I.*
